# Rhomboid protease RHBDL4/RHBDD1 cleaves SREBP-1c at ER monitoring and regulating fatty acids

**DOI:** 10.1101/2021.08.24.457590

**Authors:** Song-iee Han, Masanori Nakakuki, Yoshimi Nakagawa, Yunong Wang, Masaya Araki, Yuta Yamamoto, Hiroaki Tokiwa, Hiroyuki Takeda, Yuhei Mizunoe, Kaori Motomura, Hiroshi Ohno, Yuki Murayama, Yuichi Aita, Yoshinori Takeuchi, Yoshinori Osaki, Takafumi Miyamoto, Motohiro Sekiya, Takashi Matsuzaka, Naoya Yahagi, Hirohito Sone, Hiroyuki Kawano, Hitoshi Shimano

**Author notes:** These authors contributed equally (Song-iee Han, Masanori Nakakuki). Hitoshi Shimano, MD, Ph.D, Department of Endocrinology and Metabolism, Faculty of Medicine, University of Tsukuba, 1-1-1 Tennodai, Tsukuba, Ibaraki 305-8575, Japan.

## Abstract

The ER-embedded transcription factors, sterol-regulatory element-binding proteins (SREBPs), master regulators of lipid biosynthesis, are transported to Golgi for proteolytic activation to tune cellular cholesterol levels and regulate lipogenesis. However, mechanisms by which the cell responds to the levels of saturated or unsaturated fatty acids remain underexplored. Here we show that RHBDL4/RHBDD1, a rhomboid family protease, directly cleaves SREBP-1c at ER. The p97/VCP, AAA-ATPase complex then acts as an auxiliary segregase to extract the remaining ER-embedded fragment of SREBP-1c. Importantly, the enzymatic activity of RHBDL4 is enhanced by saturated fatty acids (SFAs), but inhibited by polyunsaturated fatty acids (PUFAs). Genetic deletion of RHBDL4 in mice fed on a Western diet enriched in SFAs and cholesterol prevented SREBP-1c from inducing genes for lipogenesis, particularly for synthesis and incorporation of PUFAs, and secretion of lipoproteins. The RHBDL4–SREBP-1c pathway reveals a regulatory system for monitoring fatty acid composition and maintaining cellular lipid homeostasis.

**Graphical Abstract:** 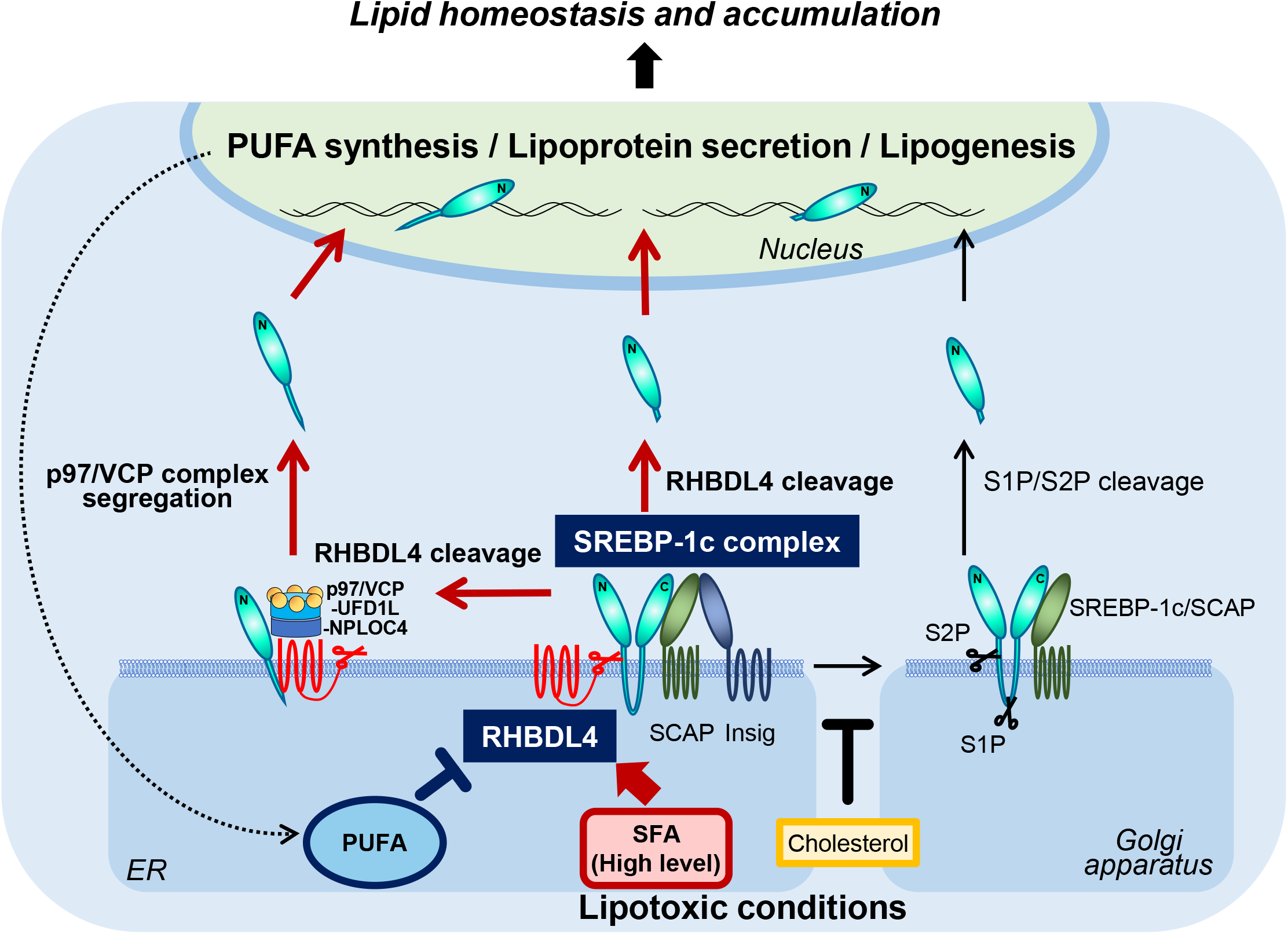

## Introduction

Sterol-regulatory element-binding protein-1a, -1c, and -2 (SREBP-1a, -1c, and -2) are a transcription factor family that regulate the biosynthesis and uptake of lipids (Horton et al., 2002; Shimano and Sato, 2017). SREBP-1a regulates the production of phospholipids, fatty acids, and cholesterol, providing the entire repertoire of membrane lipids for cell proliferation. SREBP-1c is involved in the synthesis of fatty acids and triglycerides in liver and adipose tissues. SREBP-2 acts as the master regulator to maintain cholesterol levels, controlling the expression of the full set of cholesterol synthesis-related genes and the low-density lipoprotein receptor (LDLR) gene, in response to the demands on cellular sterols (Horton et al., 1998; Shimano et al., 1997; Shimomura et al., 1997). SREBP-1c and SREBP-2 have contrasting activation mechanisms. SREBP-1c activation is mainly thought to be regulated transcriptionally, which is dependent on the intake of carbohydrates or elevated insulin. By contrast, SREBP-2 activation is regulated post-translationally through proteolytic cleavage system in cytoplasm (Horton et al., 2002; Shimano and Sato, 2017). The SREBP protein is located as a precursor form in the ER membrane and nuclear envelope. To fulfill the transcriptional role of SREBP in nucleus, it needs to depart from the ER membrane, following the proteolytic cleavage of SREBP. The proteolytic processes are mainly regulated by the levels of cholesterol and its sterol derivatives in the cell, particularly in ER. Upon sterol deprivation, SREBP precursor moves from the ER and shifts to Golgi apparatus by escort of the sterol-sensor protein, SREBP-cleavage-activating protein (SCAP). Subsequently, SREBP receives two steps of intramembrane proteolysis by site-1 protease (S1P) and site-2 protease (S2P) located in the Golgi membrane. The released N-terminal fragment of SREBP is transported to the nucleus by importin beta. If cellular cholesterol is abundant, the ER-retaining protein, insulin induced genes (Insigs) interact with the SREBP– SCAP complex and prevent their shift to the Golgi apparatus. Thus, the proteolytic cleavage of SREBP-2 is strictly regulated by this sterol-dependent system to maintain cholesterol homeostasis (Brown and Goldstein, 1999).

The proteolytic process for activation of SREBP-1c is somewhat different feature from that of SREBP-2. The SREBP-1c processing system is influenced by various factors, such as insulin, carbohydrates, and polyunsaturated fatty acids (PUFAs) in addition to sterols (Jeon and Osborne, 2012). PUFAs show obvious inhibitory effect on the proteolytic process of SREBP-1, but is much less effect on SREBP-2 (Yahagi et al., 1999). Based on the difference in selectivity of PUFA between SREBP isoforms, we hypothesized that alternative pathway for SREBP-1c activation may exist, which is independent from the proteolysis by S1P and S2P. Using mutant SREBPs in which the known cleavage sites of SREBP are disrupted by introducing alanine point mutations into the S1P, S2P, and caspase 3 (CPP32) sites, we previously showed that mutant SREBP-2 was incapable of generating the nuclear form, but that the corresponding mutant of SREBP-1c retained the potential to produce its nuclear form (Nakakuki et al., 2014). We observed that this proteolytic process of mutant SREBP-1c was still suppressed by EPA. From these differences in cleavage modes of SREBP-2 and -1c, we suspected that control of SREBP-1 may be dictated by unknown proteases sensitive to PUFAs (Nakakuki et al., 2014). Furthermore, using the antibiotic Brefeldin A as an inhibitor of the transport of ER proteins to the Golgi and various protease inhibitors as tools, it was shown that this alternative process of SREBP-1c occurs at ER by serine protease(s) probably without shifting to Golgi (Nakakuki et al., 2014).

In an attempt to seek for the responsible enzyme, we remark the rhomboid proteases from the protease group of regulated intramembrane proteolysis, in which S2P is also included (Lemberg, 2011). The rhomboid family match the conditions in that the members are intramembrane serine proteases, cleave the intramembrane helices of substrates and those activities are regulated by phospholipid environments in cell membrane (Bergbold and Lemberg, 2013; Urban and Wolfe, 2005). In human, five enzymatically-active rhomboid proteases are known; RHBDL1, RHBDL2, RHBDL3, RHBDL4, and PARL (Bergbold and Lemberg, 2013). Among them, RHBDL4 was assumed to be promising candidate that resides in ER membrane (Bergbold and Lemberg, 2013; Urban and Wolfe, 2005). In this study, we found that RHBDL4 is responsible for the proteolytic cleavage of SREBP-1c under PUFA regulation and that this pathway is involved in lipogenic genes regulated by fatty acids.

## Results

### RHBDL4 cleaves membrane SREBP-1c in a fashion independent of SCAP-Insig system

To identify the unknown enzyme responsible for PUFA-regulated cleavage of SREBP-1c, we surveyed serine proteases associated with regulated intramembrane proteolysis (RIP), based upon our previous results (Nakakuki et al., 2014). We comprehensively screened mammalian rhomboid proteases. The membrane form of human SREBP-1c tagged at the N-terminus with HSV epitope tag was cotransfected into HEK293 cells along with the five human rhomboid proteases. By western blot analysis using an antibody to the HSV tag, production of the short forms of SREBP-1c was examined in nuclear extracts to test whether any of those enzymes might cleave SREBP-1c and generate the nuclear fragment. Only RHBDL4 cleaved SREBP-1c as compared to empty vector controls (Figure 1A). Transfection of RHBDL4 also enhanced cleavage of SREBP-1a that runs at slower mobility than SREBP-1c in nuclear extracts (Figure 1B). By contrast, the effect of RHBDL4 on precursor SREBP-2 resulted in multiple bands, whereas the effect of SCAP resulted in a single band of the SREBP-2 nuclear form, consistent with the classical cleavage by S1P and S2P after escort to the Golgi (Figure S1A). The serine residue at 144 amino acid in RHBDL4 is critical for its protease activity on other substrates, such as the α chain of the pre-T cell receptor (pTα) (Fleig et al., 2012; Wunderle et al., 2016). The catalytically dead mutant, RHBDL4 (S144A) attenuated the production of cleaved SREBP-1c in nuclear extracts (Figure 1C), indicating that this residue is critical for cleavage of SREBP-1c. Consistent as the sterol-regulated SREBP-cleavage system, SCAP overexpression increased nuclear SREBP-1c protein, and additional coexpression of Insig1 suppressed the SCAP-enhanced SREBP-1c cleavage (Figure 1D). Strikingly, RHBDL4-induced SREBP-1c cleavage resulted in more pronounced accumulation of nuclear SREBP-1c compared to SCAP. RHBDL4 products from SREBP-1c were broad, containing a part migrating more slowly than that from SCAP induction, suggesting that the RHBDL4-induced cleavage of SREBP-1c may occur at multiple sites proximal to the cleavage site by S2P (Figure 1D). By contrast, Insig1 did not affect RHBDL4-induced SREBP-1c cleavage (Figure 1D). Next, we compared the effects of RHBDL4 and SCAP on a 3M mutant which entirely prevents sterol-regulated and apoptosis-induced cleavages (Nakakuki et al., 2014). The increase of nuclear SREBP-1c protein after SCAP overexpression observed for the wild type (WT) was completely abrogated by replacement with the 3M mutant SREBP-1c. Importantly, RHBDL4 exhibited similarly strong activation of cleavages for both WT and 3M mutant SREBP-1c, indicating that the cleavage by RHBDL4 occurs at different site (s) from those for S1P, S2P, and CPP32 (Figure 1E). Then, we tested whether RHBDL4 is the only requisite for SREBP-1c cleavage. Pure human recombinant SREBP-1c protein and human RHBDL4 protein were prepared using a cell-free protein expression system derived from wheat germ. These proteins were mixed in detergent micelles to evaluate the direct proteolytic activities of RHBDL4 (Figure 1F). This reconstitution of proteolysis demonstrated that RHBDL4 is able to robustly generate nuclear-sized SREBP-1c from the decreased precursor protein, thereby confirming that RHBDL4 rhomboid activity is sufficient for cleavage of SREBP-1c. Moreover, RHBDL4-induced SREBP-1cleavage observed in cultured cells was evaluated in mouse livers (Figure 1G). After adenoviral RHBDL4 infection into mice, hepatic SREBP-1 was investigated in a refed state after fasting, the condition in which endogenous precursor SREBP-1 is strongly induced. This hepatic overexpression of RHBDL4 markedly increased levels of the nuclear form of SREBP-1 protein, indicating that RHBDL4 also cleaves SREBP-1 *in vivo*.

**Figure 1.**
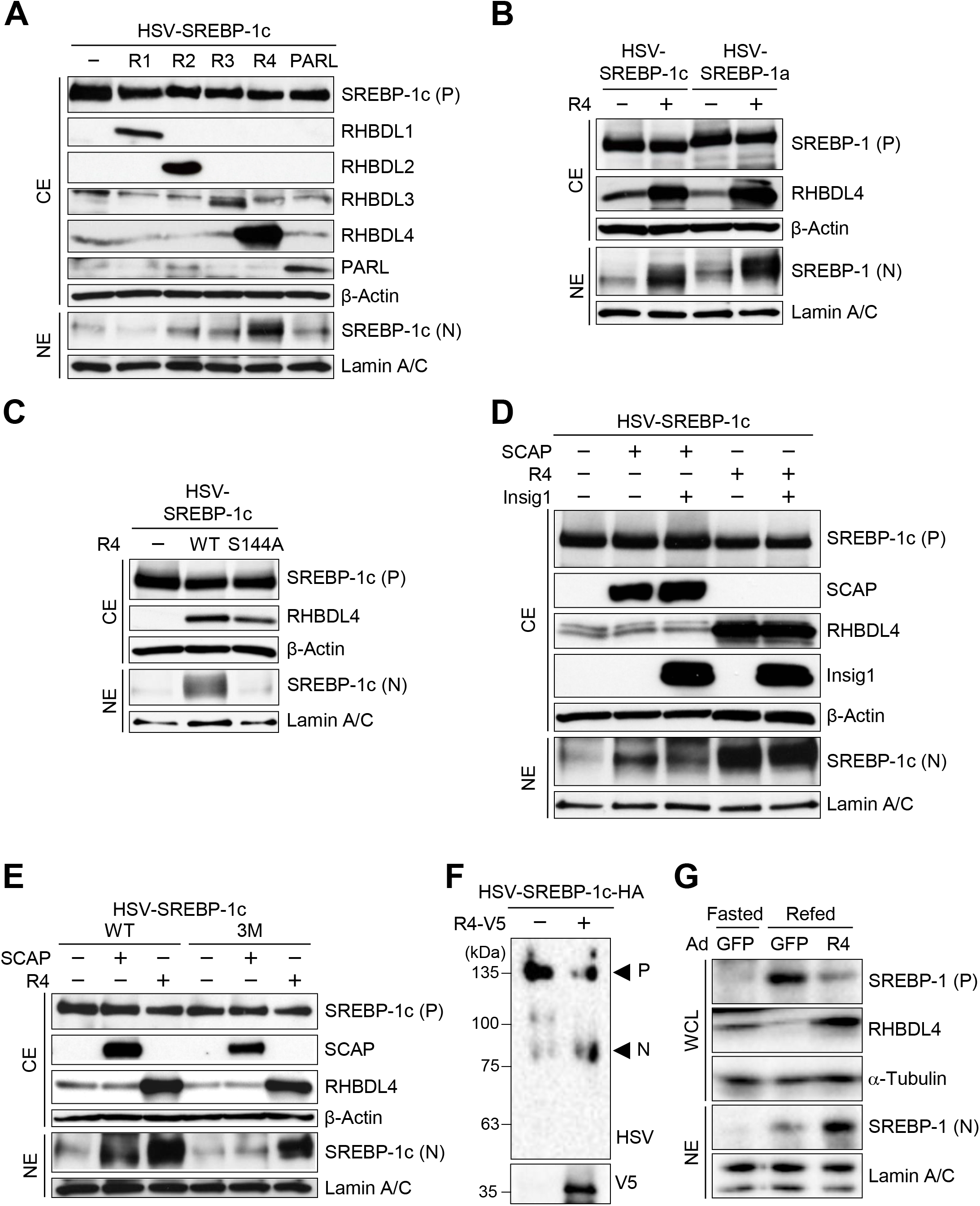
RHBDL4 cleaves SREBP-1c independently of the SCAP-Insig system. (A–E) Indicated proteins were transiently transfected into HEK293 cells. After 24 hr, cytosolic extracts (CE) and nuclear extracts (NE) were immunoblotted with the indicated antibodies. SREBP-1c cleavage by rhomboid protease family (A), SREBP-1a cleavage by RHBDL4 (B), SREBP-1c cleavage by RHBDL4 mutant S144A (C), SREBP-1c cleavage after coexpression of RHBDL4 and Insig1 (D), and SREBP-1c mutant 3M cleavage by RHBDL4 (E) are shown. HSV-SREBPs were immunoblotted with HSV antibodies. (−), mock; R1, RHBDL1; R2, RHBDL2; R3, RHBDL3; R4, RHBDL4; WT, wild type; S144A, the catalytically inactive mutant of RHBDL4; 3M, three known cleavage sites for CPP32, S2P, and S1P mutant of SREBP-1c, P, precursor; N, nuclear. (F) An *in vitro* cleavage assay was performed using recombinant HSV-SREBP-1c-HA and RHBDL4-V5 purified from a cell-free protein synthesis system using wheat germ. Indicated proteins were analyzed by immunoblotting. Arrowheads show precursor (P) and nuclear (N) SREBP-1c. (G) Eight-week-old male C57BL/6J mice were injected with green fluorescent protein (GFP) or RHBDL4 expressing adenovirus via the tail vein. Six days after injection, the mice (n = 3–4 per group) were subjected to fasting and refeeding. Immunoblot analysis of indicated proteins in whole cell lysates (WCL) and NE from mouse livers was performed. See also Figures S1 and S2.

We investigated the region containing the RHBDL4-mediated cleavage site(s) within SREBP-1c using placental alkaline phosphatase (PLAP)-SREBP-1c chimera proteins (Figures S1B and S1C). The data suggest that RHBDL4-mediated cleavage can occur at more than one site: one in the cytosol region of SREBP-1c before the first TM domain (TM1) and the other in the loop domain of SREBP-1c in the ER lumen after the TM1 (Figure S1D). Diverse prokaryotic and eukaryotic rhomboid proteases share consensus motifs: a less stringent Arg-(X)_n_-Arg motif for cleavage of substrate proteins within or the near the membrane domain (Strisovsky et al., 2009). The corresponding motif (LDRSRL) was also found in SREBP-1c, overlapping the one for S2P as depicted in Figure S1D. We made several mutants by amino acid substitution near the TM domain of SREBP-1c and tested their cleavages by RHBDL4 (Figure S1E). Consistent with PLAP data, two cleaved bands (long and short forms) from WT SREBP-1c were detected. From the mutation analysis of the LDRSRL region (Figure S1D), mutation of serine 462 to phenylalanine (S462F) prevented the shorter cleaved form of SREBP-1c, but L459F and L464F substitutions did not (Figure S1E). To specifically evaluate the effect of RHBDL4 on SREBP-1c cleavage and resultant nuclear translocation and transactivation, a reporter assay with the Gal4-VP16-SREBP-1c system was used. In this reporter system, the expression of a WT SREBP-1c fusion protein exhibited a modest basal luciferase activity that was robustly enhanced by overexpression of RHBDL4 (Figure S1F). When the RHBDL4-cleavage defective SREBP-1c mutation was introduced in this system, Gal4-VP16-SREBP-1c (S462F) completely abrogated both basal activity and diminished the induction from RHBDL4 overexpression (Figure S1F). The data indicates that the short form of RHBDL4-mediated SREBP-1c cleavage contributes to SREBP-1c transactivation and that the basal activity in HEK293 cells might reflect the level of activity of endogenous RHBDL4.

The RHBDL4–SREBP-1c complex structure was modeled using the Molecular Operating Environment (MOE) program. As shown in Figure S2, the S2P recognition and cleavage site at TM1 in SREBP-1c fits a groove near the catalytic triad site of RHBDL4 (histidine 80, serine 144, and histidine 195) well, consistent with our mutational analysis. These data suggested that the DRSR region is important for substrate recognition and the resultant cleavage by RHBDL4 to the short form of SREBP-1c, but that the recognition mode should be different between these two proteases in different organelles. This is consistent with the 3M mutant that prevents S2P cleavage still being cleaved by RHBDL4.

### RHBDL4 and SREBP-1c colocalize and interact at ER to release a nuclear form of SREBP-1c

RHBDL4 was reported to be located at ER as an intramembrane protease where membrane-bound SREBP-1c typically resides (Fleig et al., 2012). Reconfirming this, it was demonstrated that RHBDL4 colocalized with an ER-marker protein and with SREBP-1c at the ER and nuclear envelope. This indicates that SREBP-1c could be a substrate for RHBDL4 at the ER and nuclear envelope directly without additional interorganelle trafficking as is observed for the SCAP–S1P– S2P system (Figure 2A). In immunoprecipitation (IP) experiments with coexpressed HSV-tagged SREBP-1c and V5-tagged RHBDL4 in HEK293A cells, the immune-reactive V5-tagged RHBDL4 signal was detected after IP with the HSV-tag antibody, indicating the proteins can interact (Figure 2B). Other combinations of complex formation were also examined by reciprocal IP experiments of RHBDL4 as well as further IP experiments with each member of the SREBP– SCAP-Insig complex. V5-tagged RHBDL4 was also immunoprecipitated with both HA-tagged SCAP and Myc-tagged Insig1 (Figures 2C and 2D). Validation of these immunoprecipitation experiments was ensured by confirming an interaction of SCAP with both SREBP-1c and Insig1 (Figure 2E). The data indicated that RHBDL4 physically interacts with SREBP-1c, SCAP, and Insig1, suggesting the possibility that RHBDL4 can form a part of the SREBP-1c–SCAP–Insig complex at the ER, although the two systems work separately.

**Figure 2.**
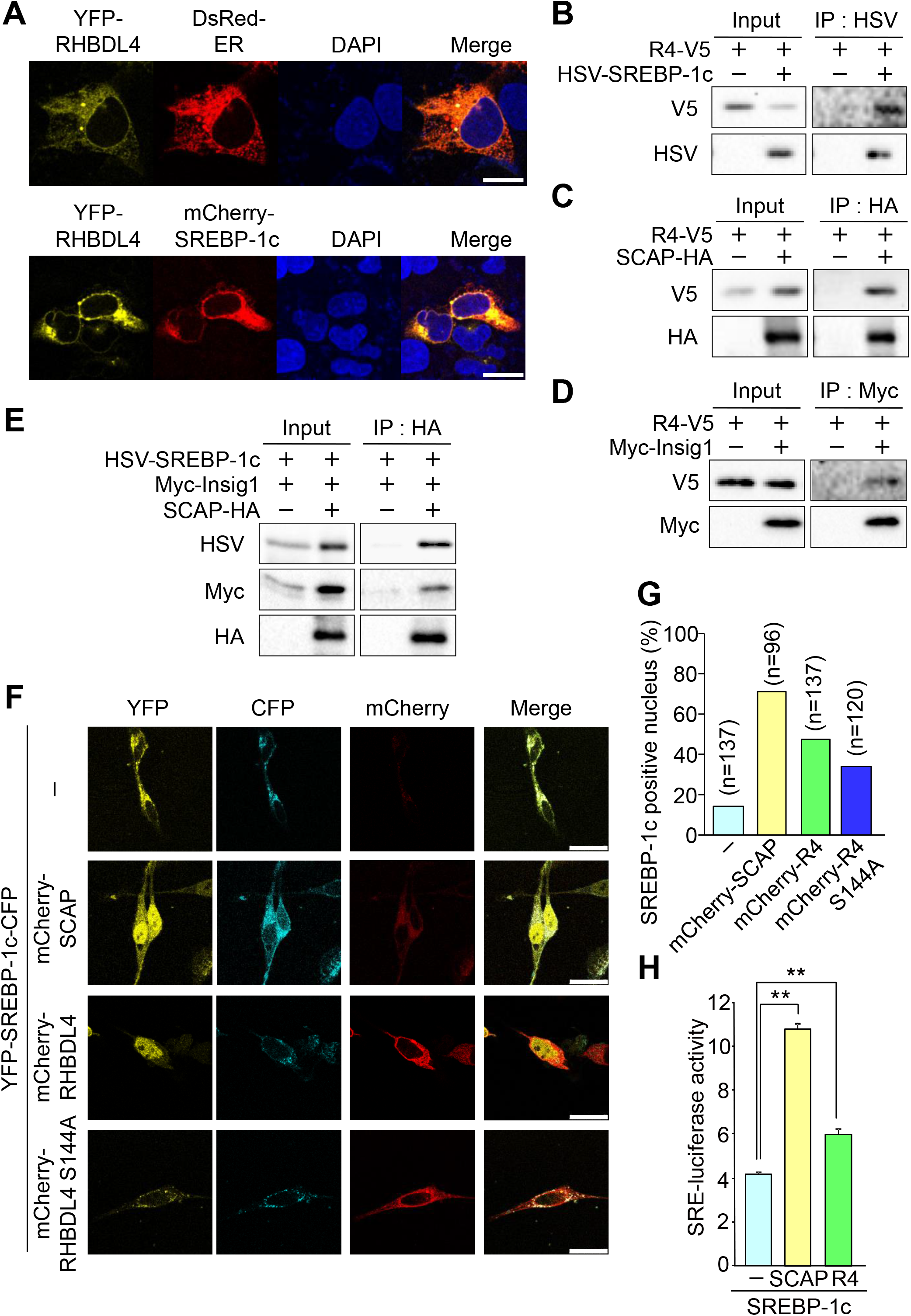
RHBDL4 interacts with the SREBP-1c–SCAP–Insig complex in ER and increases nuclear SREBP-1c accumulation. (A) Cellular localization of RHBDL4 and SREBP-1c. HEK293A cells expressing the indicated YFP-RHBDL4 and DsRed-ER (top) or mCherry-SREBP-1c (bottom) constructs are shown. Nuclei were stained with DAPI. Scale bar, 20 μm. (B–E) Immunoprecipitation assay. HEK293A cells were transfected with the indicated expression plasmids. After 24 hr, cell extracts were immunoprecipitated (IP) with the indicated antibodies. Bound proteins were immunoblotted with the indicated antibodies. Interaction of SREBP-1c and RHBDL4 (B), SCAP and RHBDL4 (C), Insig1 and RHBDL4 (D), SCAP and SREBP-1c and Insig1 (E) are shown. (F and G) Translocation of SREBP-1c to the nucleus by RHBDL4. HeLa cells expressing YFP-SREBP-1c-CFP and mCherry-SCAP, mCherry-RHBDL4, or mCherry-RHBDL4 S144A are shown. Cells with nuclear fluorescence of YFP were counted and the percentages expressed as bar graphs (G). Scale bar represents 25 μm. (H) Luciferase assay for SREBP-1c activity. HEK293 cells were transfected with SRE-Luc, pRL-SV40, and the indicated expression plasmids. After 24 hr, cell extracts were examined using luciferase assays. Quantification was performed on four samples. Data are represented as means ± SEM. ** P < 0.01.

To assess the consequence of the proteolytic cleavage of SREBP-1c by RHBDL4, the processing of SREBP-1c was visualized and translocation to the nucleus was tested. SREBP-1c doubly labeled with YFP at the N-terminus and with CFP at the C-terminus (YFP-SREBP-1c-CFP) was cotransfected with mCherry-labeled RHBDL4 or SCAP in HeLa cells, a SREBP abundant cell line. In the absence of SCAP or RHBDL4 overexpression, both YFP- and CFP-SREBP-1c signals were localized and merged in the cytosol assuming to be at ER according to the data from Figure 2F. After mCherry-tagged SCAP coexpression, the signal of N-terminal YFP-SREBP-1c shifted to the nucleus, whereas the signal of C-terminal CFP-SREBP-1c remained in the cytosol. After coexpression of mCherry-RHBDL4, the signal of N-terminal YFP-SREBP-1c similarly shifted to the nucleus, indicating transfer of N-terminal SREBP-1c into nucleus. The mCherry-RHBDL4 mutant that was observed to be catalytically defective *in vitro* mostly abrogated the nuclear appearance of SREBP-1c (Figure 2F). The quantitative data for SREBP-1c positive nuclei indicated that overexpression of RHBDL4 induced a nuclear shift of SREBP-1c to a comparable extent to SCAP, dependent on its enzymatic activity (Figure 2G).

We then tested if nuclear translocation also resulted in transactivation of SREBP-1c using a luciferase based reporter assay with a SREBP binding cis-element (SRE) in the promoter region. Coexpression of the membrane forms of SREBP-1c with either SCAP or RHBDL4 significantly elevated the transcriptional activity, although less strongly after RHBDL4 expression (Figure 2H). Taken together these data in HeLa cells indicate that RHBDL4 overexpression results in N-terminal SREBP-1c cleavage, nuclear translocation, and consequent transcriptional activity.

### RHBDL4 siRNA knockdown reduces nuclear SREBP-1 protein and suppresses target gene expression

We hypothesized from control data in our overexpression experiments that there may be a contribution from endogenous RHBDL4. To test the physiological contribution of RHBDL4 to SREBP-1 processing, RHBDL4 knockdown (KD) experiments were performed to investigate the effects on the levels of cleaved SREBP-1 protein in nuclear extracts and its target gene expression in HepG2 and HEK293 cells (Figure 3). Two independent RHBDL4 siRNAs effectively abolished endogenous RHBDL4 protein in both cell lines. We observed that the classical sterol regulation in these cells obscured the effect from endogenous RHBDL4 (Figure 3). To suppress the endogenous effect mediated from SCAP, 25-hydroxycholesterol was added to inhibit the SCAP-S1P-S2P process. HepG2 and HEK293 cells that were transfected with RHBDL4 siRNA showed a clear reduction of the amount of nuclear SREBP-1 protein without changes in the levels of the membrane form of SREBP-1 (Figures 3A and 3B). Gene expression analysis revealed RHBDL4 siRNA diminished *Rhbdl4* (*Rhbdd1*) mRNA, with a slight concomitant reduction of *Srebp-1* (*Srebf1*) mRNA in HepG2 cells. SREBP-1 target genes; stearoyl-CoA desaturase 1 (*Scd1*), fatty acid desaturase 1 (*Fads1*), and thyroid hormone responsive (*Thrsp*) were all significantly suppressed by both of the two RHBDL4 siRNAs (Figure 3C). The data indicate that endogenous RHBDL4 is involved in SREBP-1 activation and subsequent gene regulation of its target genes independently from sterol regulation. The RHBDL4 KD also reduced the gene expression of *Srebp-2* (*Srebf2*) and its target gene, *Ldlr* (Figure 3C). In contrast, *S1P* (*Mbtps1*) was upregulated, implicating both SCAP and RHBDL4 systems might be coregulated as observed in the obscure data without 25-hydroxycholesterol treatment (Figures S3B and S3C).

**Figure 3.**
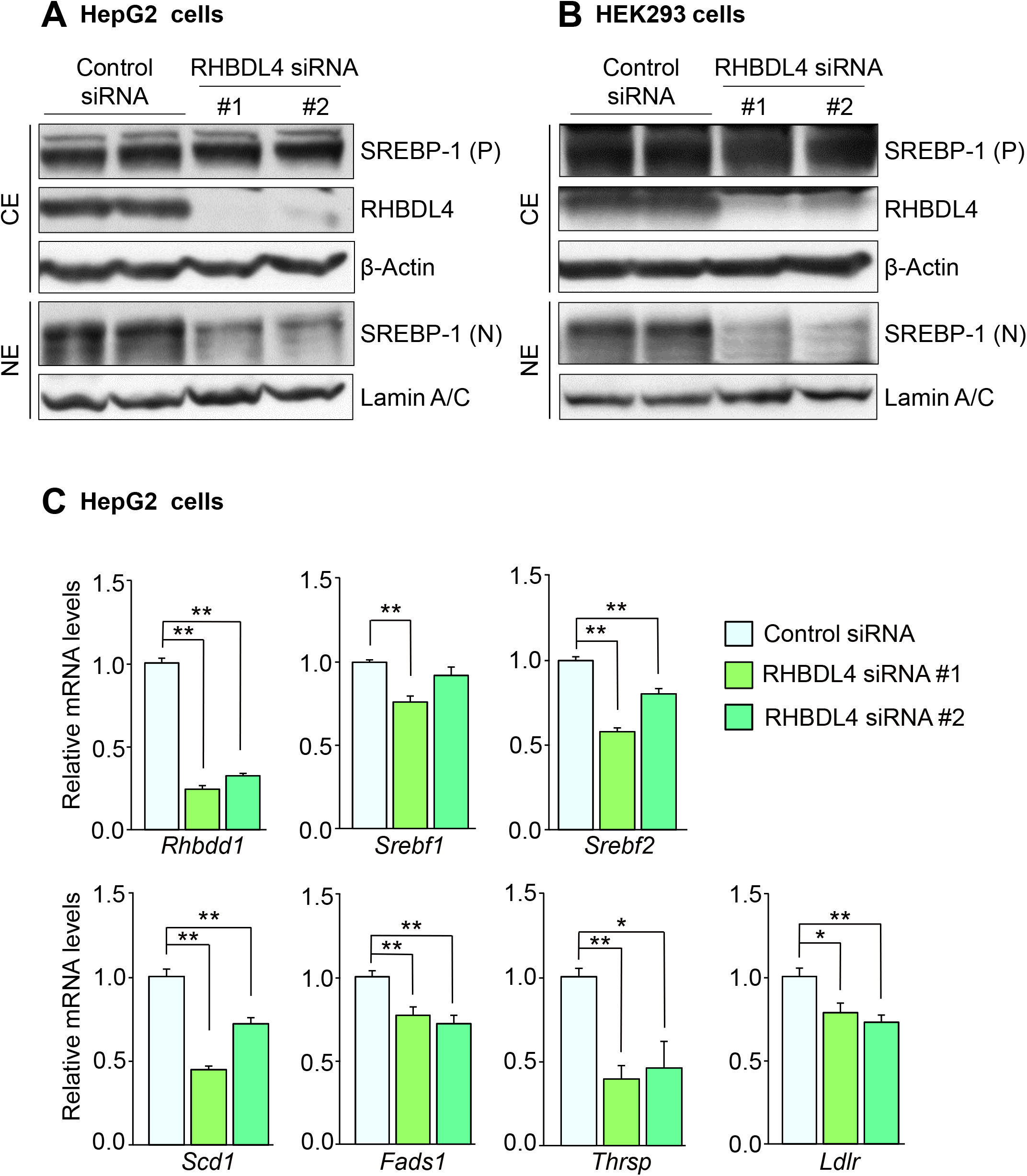
siRNA knockdown of RHBDL4 reduces nuclear SREBP-1 and suppresses target gene expression. (A–C) HepG2 cells (A and C) and HEK293 cells (B) were transfected with control siRNA or two independent RHBDL4 siRNAs (#1 and #2). Twenty-four hr after transfection, cells were incubated with 10% DLS containing 3 μM 25-hydroxycholesterol for 24 hr. (A and B) Immunoblot analysis of the indicated proteins in cytosolic extracts (CE) and nuclear extracts (NE). P, precursor; N, nuclear. (C) qRT-PCR analysis of SREBP-1 and related genes. Data are represented as means ± SEM. Quantification was performed on six samples. * P < 0.05, ** P < 0.01. See also Figure S3.

### RHBDL4 is responsible for activation of hepatic SREBP-1cleavage and lipogenic gene expression in mice on a Western diet

To clarify the role of RHBDL4 in SREBP-1cleavage in the liver, RHBDL4 knockout (KO) mice were generated using CRISPR/Cas9. RHBDL4 KO mice were alive and exhibited no apparent anthropometric phenotypes (Figures S4A and S4B). On a normal chow diet, the disappearance of hepatic RHBDL4 protein and mRNA were confirmed in the livers of RHBDL4 KO mice, but there were no marked changes in either membrane or nuclear SREBP-1 protein between WT and RHBDL4 KO mice (Figures 4A and 4B). The animals were then fed on a Western diet (WD) enriched with cholesterol and SFA for 14 days whereby the membrane form of SREBP-1 in liver were robustly induced due to the gene induction by LXR–SREBP-1c pathway, resulting in a striking increase in the nuclear form in WT mice. In spite of a similar robust increase in precursor SREBP-1 protein after feeding with a WD, the amount of SREBP-1 protein in nuclear extracts in RHBDL4 KO mice was strongly reduced. This is indicative of the suppression of SREBP-1 processing in the absence of RHBDL4. RHBDL4 appears to contribute to the cleavage of SREBP-1 in the liver as a response to feeding on a WD. Membrane and nuclear SREBP-2 proteins were not different between the WT and KO mice (Figures 4A and 4B). We also evaluated the effect of RHBDL4KO on another SREBP-1c-induced condition: a high-sucrose fat-free (HS) diet. The reduction in nuclear form SREBP-1 from RHBDL4 KO mice was also observed on a HS diet, but less prominently when compared to the effects after feeding on a WD (Figure S4C).

**Figure 4.**
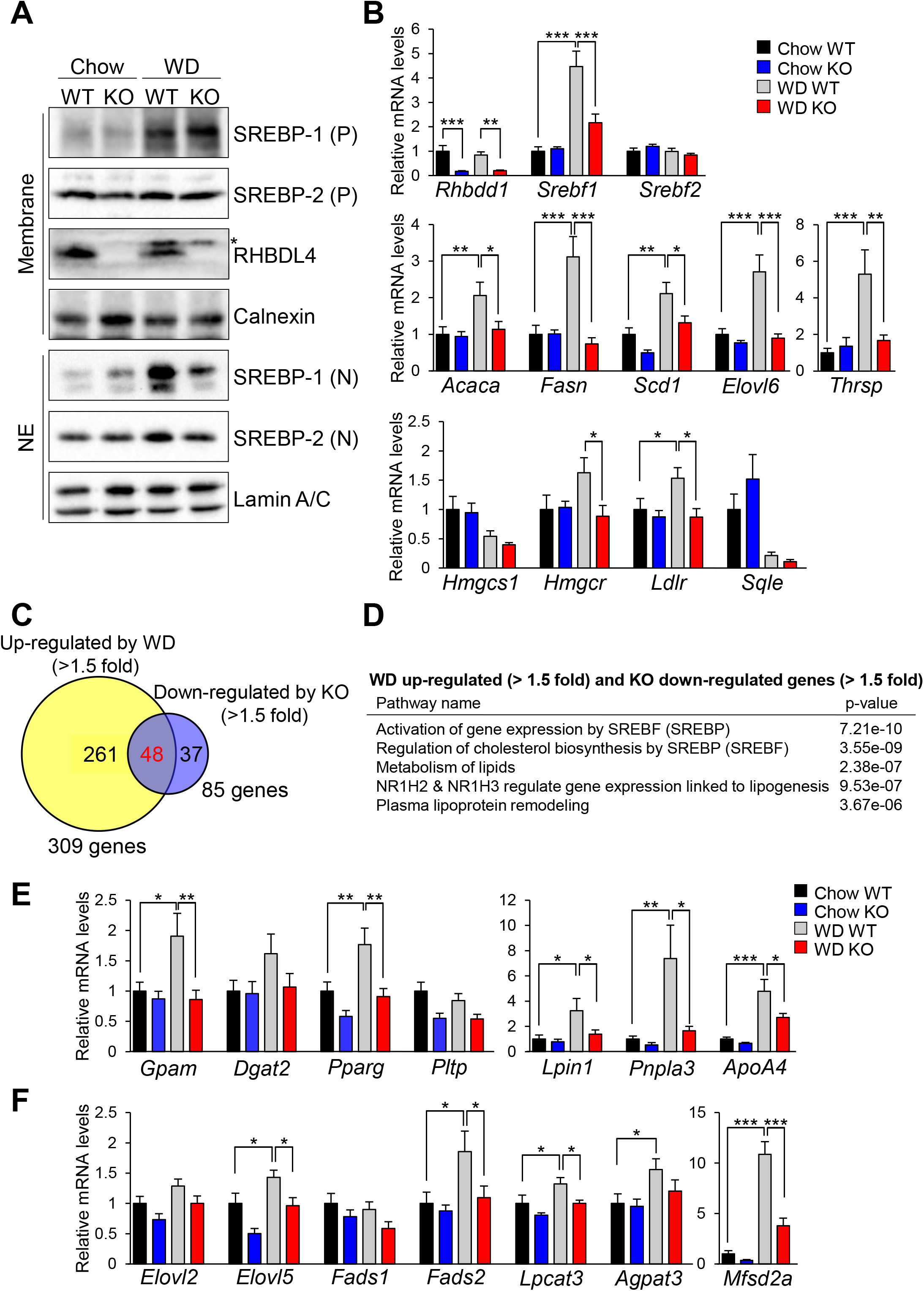
RHBDL4 contributes to SREBP-1cleavage and activation in the Western diet-fed mice. Eight-week-old male wild type (WT) and RHBDL4 knockout (KO) mice were fed on normal chow or a Western diet (WD) for 14 days. (A) Representative immunoblot analysis of the indicated proteins in the membrane fraction and nuclear extracts (NE) of mouse livers. P, precursor; N, nuclear. (B) qRT-PCR analysis of lipogenic genes from mouse livers. (C and D) RNA-seq analysis of livers of WT and KO mice fed on normal chow or a WD for 14 days (FDR < 0.05, n = 4). (C) Venn diagram of overlap between upregulated genes from mice fed on a WD (WD induced genes > 1.5-fold versus normal chow, top) and downregulated genes in KO (KO reduced genes > 1.5-fold versus WT mice fed on a WD, bottom). (D) Functional annotations associated with KO downregulated genes compared to WT mice fed on a WD and overlapped genes with WD upregulated genes compared to normal chow. (E and F) qRT-PCR analysis of lipoprotein assembly and remodeling genes (E) and PUFA producing genes (F) in mouse livers. Data are represented as means ± SEM. Quantification was performed on eight samples. * P < 0.05, ** P < 0.01 and *** P < 0.001. See also Figure S4.

qRT-PCR analysis in mouse livers determined the effects of RHBDL4 disruption on the expression of SREBP target genes (Figure 4B). *SREBP-1c* (*Srebf1*) gene expression was markedly induced by feeding on a WD, whereas *Srebp-2* (*Srebf2*) gene did not change among the four groups. Hepatic *de novo* lipogenic genes as SREBP-1 target genes, such as acetyl-CoA carboxylase alpha (*Acaca*), fatty acid synthase (*Fasn*), *Scd1*, ELOVL fatty acid elongase 6 (*Elovl6*), and *Thrsp*, were increased in WT on a WD, and markedly suppressed in RHBDL4 KO mice. These changes corresponded to the changes in amounts of nuclear SREBP-1 in the two groups. In contrast, expression of cholesterol synthetic and uptake genes, such as 3-hydroxy-3-methylglutaryl-CoA synthase 1 (*Hmgcs1*) and reductase (*Hmgcr*), *Ldlr*, and squalene epoxidase (*Sqle*) genes, were slightly affected by RHBDL4 disruption or WD. To further understand the roles of RHBDL4 in the liver from gene expression profiles, livers from RHBDL4 KO and WT mice fed on a normal chow and on a WD were subjected to RNA-seq analysis. The analysis identified 309 genes upregulated after WD feed compared with the normal chow group, and 85 genes were downregulated by RHBDL4 KO compared to WT (Figure 4C). Gene ontology analysis on the 48 overlapping genes resulted in enrichment of SREBP-regulated genes and the LXR pathway, consistent with induction of the LXR–SREBP-1c axis (Figure 4D). Thus, our identification of RHBDL4 as being responsible for SREBP-1cleavage and downstream effects on transcription was validated for having an endogenous role in the liver. It is noteworthy that SREBP-1cleavage by RHBDL4 is prominent only after feeding on a WD.

### RHBDL4 directs SREBP-1c activation to PUFA synthesis and remodeling, and VLDL secretion

The genes downregulated by RHBDL4 KO resulted in similar gene ontology enrichment, suggesting a deep relationship between RHBDL4 and SREBP in lipid metabolism, especially for plasma lipoprotein assembly and remodeling (Figure 4D). As confirmed by qRT-PCR (Figure 4E), other gene clusters upregulated after WD feeding in WT mice, but abrogated in RHBDL4 KO mice included genes involved in the incorporation of fatty acids into glycerolipids for the formation of triglycerides, such as glycerol-3-phosphate acyltransferase (*Gpam*), diacylglycerol O-acyltransferase 2 (*Dgat2*), peroxisome proliferator activated receptor gamma (*Pparg*), lipid-influx related genes, such as Lipin1 (*Lpin1*), and patatin like phospholipase domain containing protein 3 (*Pnpla3*), along with lipoprotein remodeling genes for the secretion of VLDL, such as apolipoprotein A4 (*ApoA4*) (VerHague et al., 2013) and lysophosphatidylcholine acetyltransferase 3 (*Lpcat3*) (Hashidate-Yoshida et al., 2015). Changes in these genes suggest that RHBDL4 could be involved in lipid flux between organs via circulation. Genes involved in modification or remodeling of PFA were also enriched (Figure 4F): *Elovl2* (Pauter et al., 2014), *Elovl5* (Moon et al., 2009), *Fads1*, and *Fads2* (Varin et al., 2015) for the production of long PUFA, *Lpcat3* (Rong et al., 2017), and 1-acylglycerol-3-phosphate O-acyltransferase 3 (*Agpat3*) (Hishikawa et al., 2020) for the incorporation of EPA and docosahexaenoic acid (DHA), respectively, into membrane phospholipids. Major facilitator superfamily domain containing protein 2a (*Mfsd2a*), DHA specific transporter protein showing an outstanding difference by RHBDL4 absence (Ben-Zvi et al., 2014; Chan et al., 2018; Nguyen et al., 2014). These data suggest that RHBDL4 play a role to maintain membrane PUFA levels counter-balancing lipotoxic stress of SFA and cholesterol by activating these SREBP-1 target genes through RHBDL4– SREBP-1c and LXR–SREBP-1c pathways. The findings also suggest a potential feedback system between SREBP-1 and PUFA. In terms of physiological relevance, upregulated gene clusters in RHBDL4 KO mice on a normal chow diet included ER stress and UPR related gene pathways, whereas downregulated genes related to the heat stress response and plasma lipoprotein remodeling (Figure S4D). In accordance that ER stress was reported after RHBDL4 deletion in cell culture experiments (Fleig et al., 2012), RHBDL4 might be involved in ER membrane stresses through lipid perturbations.

### RHBDL4 deletion ameliorates lipid accumulation by WD

The plasma and liver lipids, fatty acid composition, and histological analyses were shown in Figure 5A. Both genotypes after WD feeding increased liver and plasma cholesterols, and liver triglycerides. RHBDL4 KO mice showed significantly less plasma and liver triglycerides than WT mice, consistent with the differences in lipogenic and lipoprotein-related genes in the liver. Gas chromatography analysis of the fatty acid composition of liver lipids showed that RHBDL4 KO mice had decreased total fatty acid contents (Figure 5B). Most individual fatty acids including SFA and PUFA exhibited significant reduction in RHBDL4 KO livers consistent with suppressed lipogenesis. Notably, quantities of arachidonic acid (AA), EPA, docosapentaenoic acid (DPA), and DHA were decreased in RHBDL4 KO mice, consistent with decreased expression of genes required for the incorporation of AA and DHA. On a relative molar% basis, γ–linolenic acid was increased whereas palmitoleic acid and dihomo-γ–linolenic acid were decreased (Table S1). The ratio of representative ω3 PUFA to ω6 PUFA was also reduced, indicating a trend of proinflammation. Liver histology from hematoxylin and eosin (H&E) staining indicated no marked changes in RHBDL4 KO mice compared to WT on a normal chow (Figure 5C). As shown in Oil red O staining, feeding on a WD caused hepato-steatosis in both genotypes with amelioration of the size and the number of lipid droplets in RHBDL4 KO mice especially in central vein areas compared to WT (Figure 5D).

**Figure 5.**
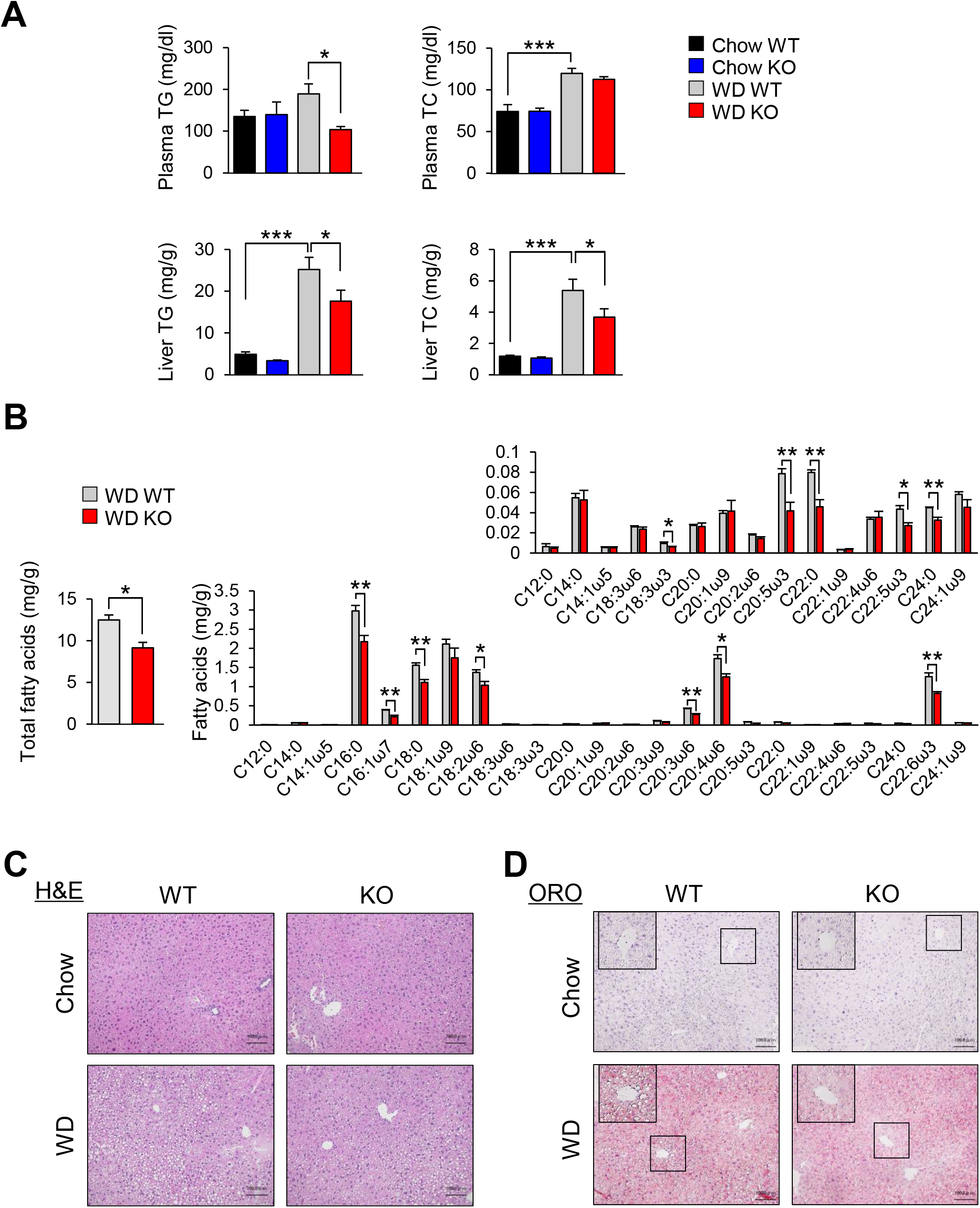
RHBDL4 deletion reduces lipid accumulation by Western diet. Eight-week-old male wild type (WT) and RHBDL4 knockout (KO) mice were fed normal chow or a Western diet (WD) for 14 days. (A) Plasma triglyceride (TG), plasma cholesterol (TC) levels, liver TG, and liver TC contents of mice. Data are represented as means ± SEM. Quantification was performed on eight samples. * P < 0.05, *** P < 0.001. (B) Total fatty acids (left) and fatty acid composition (right) of liver tissue in WT and KO fed a WD for 14 days. Data are represented as means ± SEM. Quantification was performed on four samples. * P < 0.05, ** P < 0.01. (C and D) Representative hematoxylin and eosin (H&E) stained section (C) and Oil red O (ORO) stained section (D) of mouse livers. Scale bar represents 100 μm. See also Table S1.

### PUFA inhibits and SFA activates RHBDL4-dependent SREBP-1c cleavage

It is well known that hepatic SREBP-1c and target lipogenic genes are suppressed by dietary PUFA (Sato et al., 2010; Sekiya et al., 2003; Yahagi et al., 1999), but the precise molecular mechanism is enigmatic. We investigated this in RHBDL4 KO mice (Figures 6A and 6B). One single day addition of EPA, a representative PUFA on the last day of 14 days WD feeding caused marked suppression of the nuclear SREBP-1 protein in WT livers as previously reported (Sato et al., 2010; Sekiya et al., 2003) (Figure 6). By contrast, in RHBDL4 KO mice on a WD, the nuclear SREBP-1 amount was already reduced, and only marginal further inhibition was observed with EPA. This suggested that RHBDL4 is partially responsible for EPA-regulated SREBP-1cleavage. The expression profile of SREBP-1 target genes on an EPA-containing diet is shown (Figure 6B). As previously reported, *SREBP-1c* (*Srebf1*) expression exhibited a robust reduction by EPA whereas *Srebp-2* (*Srebf2*) reduction was weak (Yahagi et al., 1999). Lipogenic genes of SREBP-1 targets, such as *Fasn, Scd1*, and *Elovl6* were similarly reduced in RHBDL4 KO mice on WD feeding, but underwent no further suppression after EPA supplementation, indicating that RHBDL4 activity on SREBP-1 was prevented by EPA feeding. Inhibition of these lipogenic genes by EPA was strongly observed in WT mice, but was diminished in RHBDL4KO mice. The data can be interpreted that a long known suppression of SREBP-1c and lipogenic genes by PUFA was at least partially attributed to RHBDL4 cleavage activity for SREBP-1c. Expression of SREBP-2 downstream target genes, such as *Hmgcs1* and *Hmgcr*, did not show changes between WT and RHBDL4 KO mice irrespective of EPA addition.

**Figure 6.**
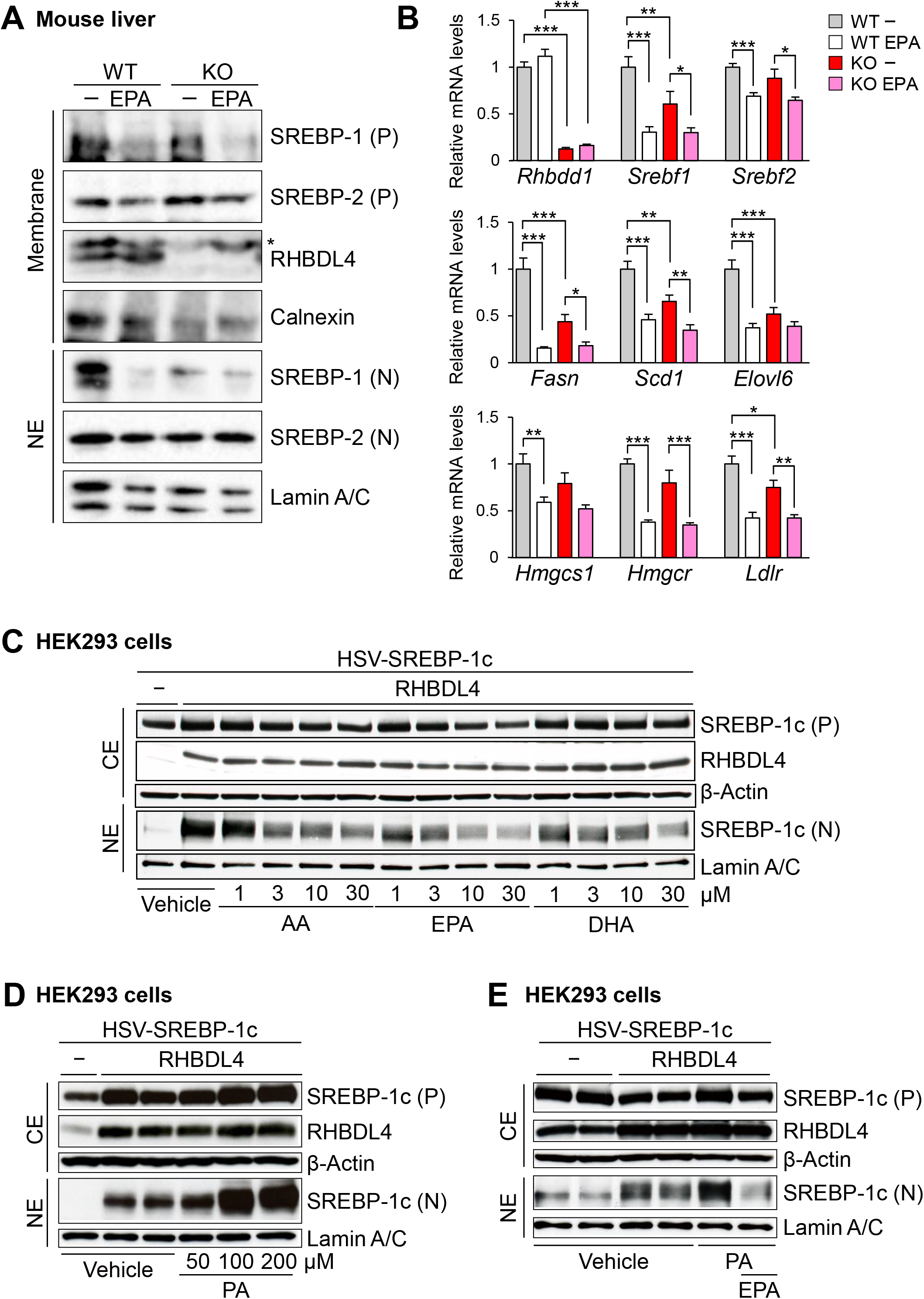
PUFA inhibits and SFA activates RHBDL4-dependent SREBP-1c cleavage. (A and B) Eight-week-old male wild type (WT) and RHBDL4 knockout (KO) mice were fed a Western diet (WD) for 14 days, and treated with 5 % EPA-E once the last day. (A) Representative immunoblot analysis of SREBP-1/2 and RHBDL4 in the membrane fraction and nuclear extracts (NE) of mouse livers. P, precursor; N, nuclear. (B) qRT-PCR analysis of lipogenic genes in mouse livers. Data represented as means ± SEM. Quantification was performed on 9-10 samples. * P < 0.05, ** P < 0.01, *** P < 0.001. (C-E) HEK293 cells were transfected with HSV-SREBP-1c and RHBDL4 expression plasmids. After 4 hr, cells were incubated with 10% DLS containing the indicated concentration of each fatty acid for 20 hr. Immunoblot analysis of the indicated proteins in cytosolic extracts (CE) and nuclear extracts (NE) was performed. SREBP-1c cleavage by AA, EPA, and DHA (C), SREBP-1a cleavage by PA (D), SREBP-1c cleavage by 100 μM PA and 10 μM EPA (E) were shown. P, precursor; N, nuclear. See also Figures S5.

*In vivo*, dietary PUFA suppresses hepatic SREBP-1c mRNA expression, and reduces the precursor protein levels. This confounds any attempt to estimate PUFA inhibition on SREBP-1cleavage directly. Instead, we investigated the effect of supplementing various fatty acids to the medium at different concentrations on RHBDL4-dependent SREBP-1c cleavage in HEK293 cells by western blot analysis under fixed overexpression of both SREBP-1c and RHBDL4. Figure 6C shows that AA, EPA, and DHA representative PUFAs all inhibit the induction of cleaved SREBP-1c by RHBDL4 in a dose-dependent manner without affecting the level of RHBDL4 protein. Effects of other various fatty acids were also tested (Figures S5A and S5B). Most unsaturated fatty acids showed dose-dependent inhibition whereas palmitic acid (PA) as a representative SFA did not inhibit at these tested concentrations. Immunoprecipitation analysis exhibited that EPA treatment did not affect the interaction between RHBDL4 and SREBP-1c, suggesting that EPA did not affect accessibility of RHBDL4 to SREBP-1c (Figure S5C). We also investigated effect of EPA on pTα and myelin protein Z (MPZ) L170R that were reported to be cleaved by RHBDL4 (Fleig et al., 2012). EPA also exhibited suppressive effects on RHBDL4 cleavage activities for these known substrates, which suggested EPA inhibition of RHBDL4 activity is not specific to SREBP-1c (Figures S5D and S5E). In contrast to no effect at lower concentrations in Figure S5A, PA at the concentrations of 50 μM or higher showed marked dose-dependent escalation of the RHBDL4-induced cleavage of SREBP-1c (Figure 6D). EPA (10 μM) efficiently reversed the agonistic effect of PA (100 μM) (Figure 6E). The balance of fatty acid saturation potentially presumably at the ER membrane is likely to be an important factor for the regulation of RHBDL4 activity. RHBDL4 may thereby be involved in ER membrane stresses through sensing and regulating the degree of fatty acid saturation.

### RHBDL4-mediated SREBP-1c activation engages p97/VCP–UFD1L–NPLOC4 complex

RHBDL4 has been reported to be involved in the ubiquitin-dependent ERAD of single-spanning and polytopic membrane proteins together with the ternary complex containing p97/valosin containing protein (VCP), UFD1L, and NPLOC4 (Greenblatt et al., 2012; Rape et al., 2001). This complex might involve dislocation of RHBDL4 substrates from the membrane into cytosol after cleavage. To evaluate whether these factors affect RHBDL4-mediated SREBP-1c activity, the Gal4-VP16-SREBP-1c system reporter assay was used. The addition of p97/VCP, UFD1L, and/or NPLOC4 additively enhanced luciferase activity as compared to RHBDL4 alone (Figures 7A and 7C). It should be noticed that the auxiliary effects of these factors were negligible without RHBDL4, indicating a functional association with RHBDL4-mediated SREBP activation (Figure 7B). Other potential modifying factors were also tested: Ubxd8 as a PUFA acceptor and UBA domain containing 2 (UBAC2) that belongs to the pseudo-rhomboid family lacking protease activity (Lee et al., 2008; Olzmann et al., 2013). Although Ubxd8 did not affect SREBP-1c cleavage activity, UBAC2 robustly inhibited RHBDL4–p97/VCP-mediated SREBP-1c activation (Figure 7D). Inhibition of RHBDL4-dependent SREBP-1c cleavage by PUFA shown in Figure 6 was completely recaptured in this reporter assay (Figure 7E). EPA suppression was minimal in the absence of RHBDL4, supporting the hypothesis that PUFA mediate the regulation of SREBP-1c cleavage via RHBDL4 (Figure 7E).

**Figure 7.**
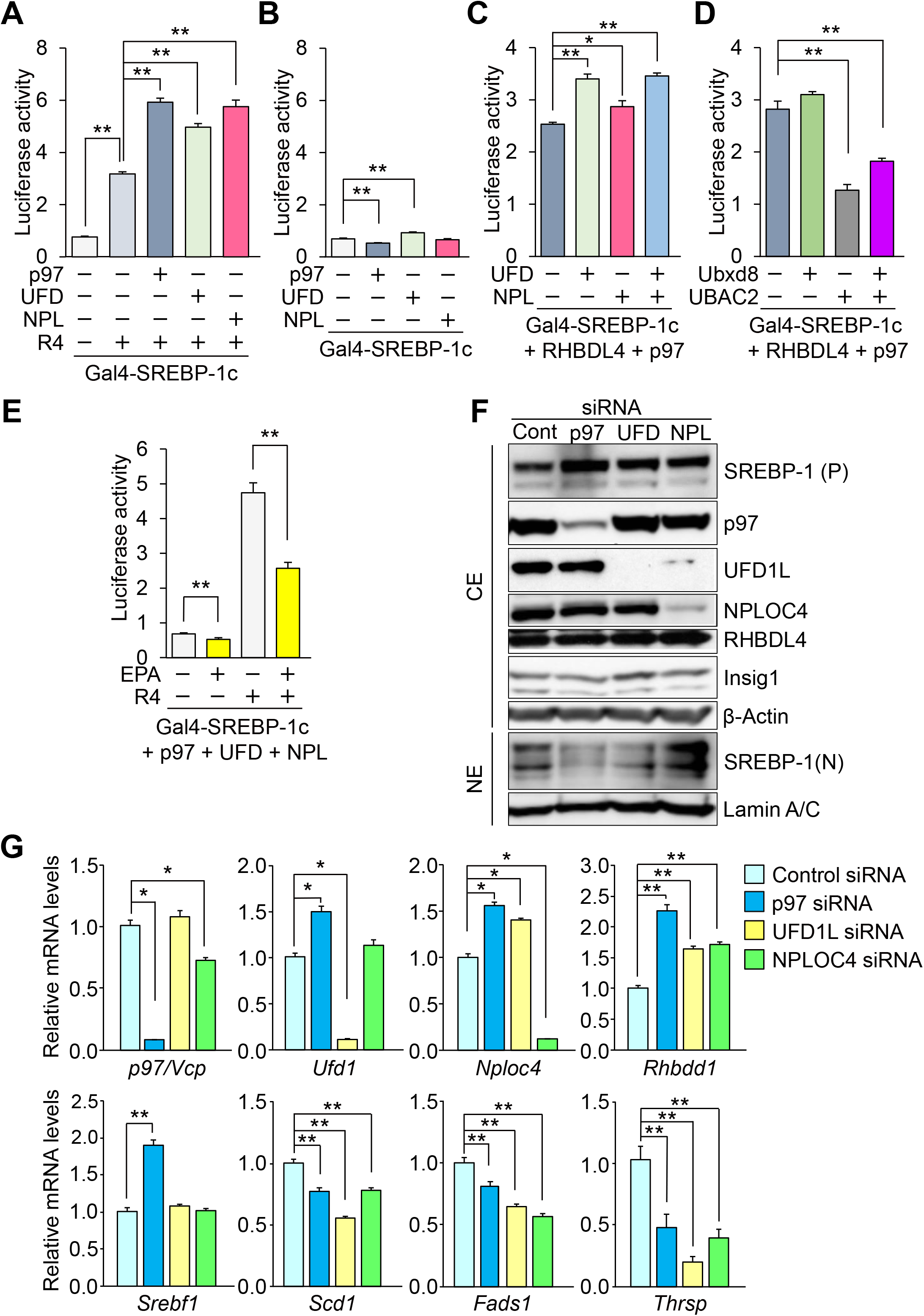
AAA-ATPase p97/VCP supports RHBDL4 induced SREBP-1c activation. (A–D) Luciferase assay of SREBP-1c activity by AAA-ATPase p97/VCP. HEK293 cells were transfected with Gal4-RE-Luc, pRL-SV40, and the indicated expression plasmids. After transfection, cells were treated with 50 μM EPA (E). After 24 hr, cell extracts were examined using luciferase assays. Quantification was performed on four samples. (F and G) HepG2 cells were transfected with the indicated siRNA. After 24 hr transfection, cells were incubated with 10% DLS containing 3 μM 25-hydroxycholesterol for 24 hr. (F) Immunoblot analysis of the indicated proteins in cytosolic extracts (CE) and nuclear extracts (NE). Cont, control; P, precursor; N, nuclear. (G) qRT-PCR analysis of SREBP-1 and related genes was performed. Quantification was performed on six samples. Data are represented as means ± SEM. * P < 0.05, ** P < 0.01. See also Figure S6.

To explore the function of p97/VCP, UFD1L, and NPLOC4 on an endogenous basis, individual siRNAs were used in HepG2 cells. Effective knocking-down by each of the three factor siRNAs was confirmed at both mRNA and protein levels (Figures 7F and 7G). NPLOC4 siRNA was found to suppress UFD1L protein level completely without affecting its mRNA level, suggesting that NPLOC4 is required for UFD1L stability (Figure 7F). Results showed that SREBP-1 protein in nuclear extracts from RHBDL4 activity was considerably reduced by p97/VCP siRNA and UFD1L siRNA, but not by NPLOC4 siRNA. Membrane SREBP-1 and RHBDL4 proteins detected in the cytosolic fraction were not changed by any of the tested siRNAs, indicating the effects on SREBP-1cleavage. As for the effects on SREBP-1 target genes, p97/VCP and UFD1L siRNAs caused a decrease in *Scd1, Fads1*, and *Thrsp* (Figure 7G). These data indicate that p97/VCP and UFD1L contributed to RHBDL4-mediated SREBP-1 activation. p97/VCP siRNA induced *Srebp-1* (*Srebf1*) expression (Figure 7G) whereas p97/VCP, UFD1L, and NPLOC4 siRNAs induced *Rhbdl4* (*Rhbdd1*), *S1P* (*Mbtps1*), and *S2P* (*Mbtps2*) expression and suppressed *Insig1* and *Insig2* expression, further supporting adaptive regulation between the RHBDL4 system and SCAP system (Figure S6A).

An auxiliary role of p97/VCP was investigated in the context of short and long forms of RHBDL-4-mediated SREBP-1c cleavage and activation (Figure S6B). The decreased, but still remnant activity in S462F SREBP-1c by RHBDL4 (Figure S1F) was enhanced by p97/VCP to a similar extent to wild SREBP-1c whereas the decrease by S462F was not changed by p97/VCP, indicating that p97/VCP enhancement is specific to the longer membrane embedded form. These data suggest that the effect of p97/VCP on RHDBL4 cleavage might be related to its segregase activity in extracting the longer cleaved SREBP-1c out of ER membranes. Roles of the versatile factor p97/VCP in lipid metabolism are not fully understood and are the target of future studies. ERAD is likely to be involved, since RHBDL4 that carry a mutation of the ubiquitin interaction motif show decreased cleavage activity for SREBP-1c (Figure S6C).

## Discussion

The current study demonstrated that rhomboid protease RHBDL4, a membrane-bound serine protease, directly cleaves SREBP-1c at the ER. This is consistent with previous data indicating that Rbd2, the Golgi yeast rhomboid homolog, cleaves Sre1, the SREBP homolog in yeast that is important for the response to hypoxia at the Golgi (Hwang et al., 2016; Kim et al., 2015). Our data highlight that RHBDL4 at the ER or nuclear envelope is regulated by fatty acids. PUFA and monounsaturated fatty acid (MUFA) are inhibitory, whereas SFA act to enhance RHBDL4 activity. In a series of both *in vitro* and *in vivo* experiments, we confirmed that RHBDL4 is responsible for a PUFA-regulated SREBP-1c cleavage system and comprise a hepatic SREBP-1c-lipogenesis pathway that is regulated by feeding on a WD and by dietary PUFA.

The process for the cleavage and transactivation of SREBP, especially SREBP-2, is tightly controlled. This occurs mediated through the cholesterol-sensing complex of SREBP2-SCAP-Insigs on the ER as the main machinery for cholesterol feedback regulation, which has been intensively analyzed and revealed by the Goldstein and Brown groups (Horton et al., 2002). In contrast, SREBP-1c is involved in lipogenesis regulated nutritionally. Nuclear SREBP-1c activity is strongly enhanced by cholesterol and SFA intakes (Repa et al., 2000; Yoshikawa et al., 2001). However, SREBP-1c is also supposed to bind to SCAP–Insigs, and thus, cannot transfer to Golgi apparatus for cleavage by S1P and S2P. In this study, we elucidated this paradox by RHBDL4 (Graphic Abstract). Retaining at ER in cholesterol abundance, N-terminal domain of SREBP-1c can be then cleaved by RHBDL4 activated in SFA abundance for nuclear transport. RHBDL4 prefers SREBP-1c-mediated lipogenesis to SREBP-2-mediated cholesterol synthesis. Detailed distinctive roles of RHBDL4 on the two isoforms of SREBPs regulated by PUFA, SFA, and cholesterol along with mutual interaction with SCAP–Insig–SREBP system are a future consideration.

Our data indicate that there are different (short and long) fragments after RHBDL4-cleavage. Once cleaved within the cytosol, the short form can be directly released into nucleus (Graphic Abstract). Another cleaved product (long fragment) is likely to be still embedded in the ER membrane and require another step for nuclear translocation (Graphic Abstract). We propose that the p97/VCP–UFD1L–NPLOC4 complex facilitates this process by extracting the ER membrane-stacking fragment. p97/VCP can exhibit this unfoldase/segregase using intrinsic ATPase (Blythe et al., 2017; Singh et al., 2019; Torrecilla et al., 2017; Xia et al., 2016). A previous study on RHBDL4 and p97/VCP suggests that their direct interaction involves ERAD (Fleig et al., 2012). By contrast, the RHBDL4 system we observed here was activating the substrate function of SREBP-1c like Nrf1 (Radhakrishnan et al., 2014), implicating p97/VCP–RHBDL4–SREBP-1c pathway as a ROS countermeasure. As a supporting evidence for the involvement of p97/VCP in this context, a p97/VCP mutation knock-in ameliorated hepatosteatosis on high fat fed mice due to impaired SREBP-1c activation (A paper in submission and personal communication from Ebihara K, Shibuya K, and Ishibashi S, Jichi Medical University).

That the protease activity of RHBDL4 varies according to the presence of different fatty acids highlights its role as a potential sensor of fatty acid saturation or possibly a sensor of the composition of the ER membrane. RHBDL4 cleavage activity on SREBP-1c was downregulated by MUFA and PUFA, but upregulated in the presence of a relatively high amount of SFA. This provides an explanation for the long known regulation of SREBP-1c by UFA. However, clear trends that were dependent on fatty acid chain length or degree of unsaturation did not emerge (Figures S5A and S5B). As a novel membrane protease feature of a rhomboid family conferred by membrane-immersion, intrinsic transmembrane segment dynamics of the rhomboid-substrate complex are thought to be important depending upon saturation of membrane fatty acids (Moin and Urban, 2012). Accordingly, our data imply that the RHBDL4–SREBP-1c interaction is preferable in the context of high SFA over PUFA and potentially relies on the ER membrane to form stable TM helices. It also needs to test the possibility of RHBDL4 as a direct acceptor or sensor for fatty acids. Further studies are needed to establish structure function relationships. Taken together with knowledge of the unique features of rhomboid family proteins in possessing a high efficiency at lateral diffusion (Kreutzberger et al., 2019), the current findings of RHBDL4 sensing SFA/UFA makes it tempting to speculate that potential allostatic responses to the RHBDL4–SREBP-1c pathway may be linked to the regulation of ER membrane fluidity.

The bona fide function of mammalian SREBP-1c has been recognized as regulating lipogenesis: the production of fatty acids and triglycerides for the supply of membrane lipids and energy storage, independent or dependent on cellular cholesterol levels. The current data in this study support RHBDL4 is a protease regulating this conventional role of SREBP-1c. Furthermore, RHBDL4 is deeply involved in regulation of PUFA metabolism (transport, synthesis, uptake and remodeling of phospholipids) in the cell membrane. Taken together with the reciprocal regulation by PUFA and SFA, RHBDL4 might be a sensing and effector of fine-tuning machinery to monitor and maintain fatty acid composition in ER membrane along with lipogenesis (Figure 4F and Graphic Abstract). In addition, according to the gene expression profile from of RHBDL4 KO mice, RHBDL4 seems to regulate ER stress-related genes. From a pathophysiological standpoint, ω-3 PUFAs possess anti-lipotoxicity and anti-inflammatory actions, protecting from fatty liver, insulin resistance, and anti-inflammatory action. The RHBDL4–SREBP-1c pathway could contribute to protect from the external stress by excessive SFA and cholesterol intakes and various inflammatory stimuli.

RNA-seq ontology analysis of RHBDL4 KO mice indicated that RHBDL4 activity is related to cellular responses to hypoxia, ROS, and fibrosis in addition to ER stress. We speculate that these regulation may be mediated by ER membrane fatty acid unsaturation both through SREBP-1-dependent and -independent pathways through other substrates of RHBDL4. We are currently evaluating chronic phenotypes of RHBDL4 KO mice after WD feeding to ascertain its pathological and clinical relevance. In the future, it will be needed to evaluate the role of RHBDL4 for lipid metabolism independent on SREBP-1c.

## Acknowledgments

We are grateful to Drs. Ken Ebihara, Daisuke Hishikawa, and Takao Shimizu for their helpful discussion. We also thank Kae Kumagai, Katsuko Okubo, Chizuko Fukui, Kenta Takei, and Takuya Kikuchi for technical assistance and the members of our laboratories for discussion and helpful comments on the manuscript. This work was supported by AMED-CREST from Japan Agency for Medical Research and Development, AMED (to H.S.); Grants-in-Aid for Scientific Research (A) 15H02541 and 18H04051 (to H.S.) and Scientific Research (C) 16K01811 and 19K11737 (to S-I.H.) from the Ministry of Science, Education, Culture, and Technology of Japan; Ono Medical Research Foundation (to S-I.H.).

The authors would like to thank Enago (www.enago.jp) for the English language review.

## Author contributions

S-I.H., M.N., Y.N., and H.Shimano designed the experiments and wrote the manuscript. S-I.H., M.N., Y.W., and M.A. performed the experiments. Y.Y. and H.Tokiwa performed computer simulation of RHBDL4–SREBP-1c complex. S-I.H., and M.N. analyzed and interpreted the data. H.Takeda, Y.Mizunoe, K.M., H.O., Y.Murayama, Y.A., Y.T., Y.O., T.Miyamoto, M.S., T.Matsuzaka, N.Y., H.Sone, and H.K. were involved in project planning and the discussion.

## STAR Methods

### RESOURCE AVAILABILITY

#### Lead Contact

Further information and requests for resources and reagents should be directed to and will be fulfilled by the Lead Contact, Hitoshi Shimano.

#### Material Availability

Plasmids and mouse lines generated in this study will be made available by request to the lead contact.

#### Data and Code Availability

RNA-seq data generated during this study is available at DDBJ database: DRA012520.

### EXPERIMENTAL MODEL AND SUBJECT DETAILS

#### Cell lines

HEK293 cells and HepG2 cells were cultured in low glucose DMEM supplemented with 10% FBS and 100 units/ml penicillin and 100 µg/ml streptomycin sulfate. HEK293A and HeLa cells were cultured in high glucose DMEM supplemented with 5% CCS or 10% FBS, respectively. All cells were cultured at 37°C in a 5% CO_2_ incubator.

#### Mice

Eight-week-old male C57BL/6J (WT) mice were obtained from CLEA Japan. For Western diet (WD) analysis, 8-week-old male mice were fed for 14 days and sacrificed in a fed state. The WD consisted of 21% fat, 20% protein, and 50% carbohydrates (Research diets, D12079B). For EPA-E treatment experiments, mice were fed on a WD for 14 days, and treated with 5% EPA-E once a last day. For high-sucrose diet (HS) analysis, 8-week-old male mice were fed for 7 days and sacrificed in a fed state. HS consisted of 70% sucrose, 10% starch, and 20% casein supplemented with methionine, vitamins, and minerals (Oriental Yeast). All animal husbandry procedures and animal experiments were performed in accordance with the Regulation of Animal Experiments of the University of Tsukuba and were approved by the Animal Experiment Committee, University of Tsukuba.

#### RHBDL4 knockout mice

Generation of RHBDL4 knockout (KO) mice using a CRISPR/Cas9 system: The px330 vector (Addgene plasmid 42230) was a gift from Dr. Feng Zhang (Cong et al., 2013). The construct including green fluorescent protein (GFP)-stop codon was inserted at the end of Exon1 of the *Rhbdl4* (*Rhbdd1*) gene, generating RHBDL4 KO. RHBDL4 CRISPR F (5′-caccGTGTATGTGTGCCATAGCACAGG-3′) and RHBDL4 CRISPR R (5′-aaacCCTGTGCTATGGCACACATACAC-3′) oligo DNAs were annealed using standard methods. Annealed DNA was purified by ethanol precipitation. Short double-stranded DNA fragments were inserted into the BbsI restriction site of the px330 vector. The constructed plasmid was designated px330-RHBDL4. Female C57BL/6J mice were injected with one shot of pregnant mare serum gonadotropin and one shot of human chorionic gonadotropin at 48-h between injections and mated with male C57BL/6J mice. Fertilized one-cell embryos were collected from oviducts. Then, 5 ng/μL of px330 vector (circular) and 10 ng/μL dsDNA donors were injected into the pronuclei of one-cell-stage embryos according to standard protocols (Gordon and Ruddle, 1981). Injected one-cell embryos were then transferred into pseudopregnant ICR mice.

## METHOD DETAILS

### Cell transfection

Cells were transfected with indicated plasmids using Fu-Gene6 transfection reagents (Roche Diagnostics), Lipofectamine 2000 transfection reagents (Thermo Fisher Scientific), or Fugene HD transfection reagent (Promega) according to the manufacturer’s instructions, and after 4 hr medium was replaced with 10% delipidated serum (DLS). DLS was prepared from FBS according to the procedure in Hannah et. al. (Hannah et al., 2001) or purchased from BioWest.

For siRNA transfection, HEK293 cells and HepG2 cells were reverse-transfected with small interfering RNAs (siRNA) in a final concentration of 33 nM using Lipofectamine RNAiMAX Transfection Reagent (Thermo Fisher Scientific). The siRNA targeted to human RHBDL4 (s38698 and s38699), human p97/VCP (s14767), human UFD1L (s14635), human NPLOC4 (s31211), and control siRNA (*Silencer*^™^ Negative Control No. 1 siRNA) were purchased from Thermo Fisher Scientific. At 1 day after transfection, medium was replaced with fresh DMEM supplemented with 10% FBS.

### Plasmids

HSV-tagged human SREBP-1c (NM_001321096), -1a (NM_004176), and SREBP-2 (NM_004599) expression plasmids were prepared by subcloning PCR products into a mammalian expression plasmid pCI (Promega). Likewise, human RHBDL1 (NM_001318733), RHBDL2 (NM_017821), RHBDL3 (NM_138328), RHBDL4 (NM_032276), PARL (NM_018622), Ubxd8/FAF2 (NM_014613), UBAC2 (NM_001144072), HA-tagged α chain of the pre-T cell receptor (pTα, BC069336), and myelin protein zero (MPZ) L170R (NM_000530.6) plasmids were prepared by subcloning DNA fragments to encode each protein into pCI. HSV-SREBP-1c point mutants were prepared using the Q5 Site-Directed mutagenesis kit (NEB). Two RHBDL4 mutants were produced by PCR using primers containing mutation according to references (Fleig et al., 2012; Wunderle et al., 2016). These mutant plasmids were firstly RHBDL4 (S144A) where protease activity is defective, and secondly RHBDL4 (ΔUIM) where the ubiquitin interacting motif was deleted by replacing leucine 274 to alanine and leucine 278 to alanine. For the *in vitro* cleavage assay, pEU-E01-MCS-HSV-human SREBP-1c-HA and pEU-E01-MCS-human RHBDL4-V5 were prepared by subcloning PCR products into the pEU-E01-MCS vector. pEU-E01-MCS vector was kindly gifted from Dr. Hiroyuki Takeda (Ehime University Proteo-Science center). mCherry-tagged human SREBP-1c, mCherry-tagged hamster SCAP (NM_001244036), mCherry-tagged human RHBDL4, mCherry-tagged human RHBDL4 (S144A), V5-tagged human RHBDL4, and Myc-tagged human Insig1 (NM_005542) were prepared by subcloning PCR products into the pcDNA3 vector (Invitrogen). YFP-tagged human RHBDL4 and YFP-human SREBP-1c-CFP were prepared by subcloning PCR products into the pCI-EYFP vector. HA-tagged hamster SCAP was prepared by subcloning PCR products into the pcDNA5/FRT/TO vector (Invitrogen). DsRed-ER plasmids were purchased from Addgene (#55836). The preparation of SCAP, Insig, SREBP-1c 3M mutant, placental alkaline phosphatase (PLAP)-human SREBP-1c (431-1123) and PLAP-human SREBP-1c (490-1123) were described previously (Nakakuki et al., 2014). Gal4-VP16-human SREBP-1c plasmid was prepared by insertion of a VP16-transactivation domain fused to a human SREBP-1c fragment from the 431st amino acid to the C-terminus (amino acids 431-1123) downstream of the Gal4-DNA binding domain sequence in pM vector (Clontech). The Gal4-RE-Luc plasmid and luciferase plasmid including sterol response element (SRE-Luc) have been described previously (Sekiya et al., 2007; Takeuchi et al., 2010). Human p97/VCP (NM_007126, SC125280), human UFD1L (NM_005659, SC320168), and human NPLOC4 (NM_017921, SC113845) expression plasmids were purchased from OriGene.

### Protein extraction and western blot analysis

Cells were treated with calpain inhibitor I directly to the medium at a final concentration of 25 µg/mL 2-h before harvest. The cytosolic and nuclear extracts were prepared using NE-PER nuclear and cytoplasmic extraction reagents (Thermo Fisher Scientific) according to the manufacturer’s protocol. Nuclear extracts and membrane proteins from mouse liver were prepared as previously described (Sheng et al., 1995). Aliquots of the proteins from cytosolic and nuclear extracts were separated by SDS polyacrylamide gel electrophoresis and transferred to PVDF membranes. The membranes were incubated with the antibodies as follows: antibodies for HSV tag (Millipore, 69171), SREBP-1 (Santa Cruz Biotechnology, sc-8984 or LifeSpan Biosciences, LS-C179707 or Novus, NB 600-582), RHBDL1 (Abnova, H00009028-A01), RHBDL2 (Abcam, ab179726), RHBDL3 (Sigma-Aldrich, SAB1104094), RHBDL4 (Proteintech, 20869-1-AP), PARL (Abnova, PAB13061), SCAP (Santa Cruz Biotechnology, sc-13553), p97/VCP (Cell Signaling Technology, #2648), UFD1L (Cell Signaling Technology, #13789), NPLOC4 (Cell Signaling Technology, #13489), Insig1 (Abcam, ab112248), V5 tag (BioLegend, #903801), HA tag (Roche, #12013819001 or Cell Signaling Technology, #3724), Myc tag (Santa Cruz Biotechnology, sc-789, sc-40, or Cell Signaling Technology, #2272), SREBP2 (Cayman chemical, #10007663), respectively. RHBDL4 antibody for mouse was constructed by Sigma-Aldrich. Antibodies for Lamin A/C (Cell Signaling Technology, #2032) as nuclear extract control, anti-α-tubulin (Sigma-Aldrich, CP06) or anti-β-actin (Cell Signaling Technology, #4970) as cytosolic control, and anti-Calnexin (BD Biosciences, #610523) were used as membrane protein control, respectively. Bound antibodies were visualized with horseradish peroxidase-conjugated IgG, ECL hyperfilm (Amersham Biosciences), and the ECL Western Blotting Analysis System or ECL Prime Western Blotting Detection Reagent according to the manufacturer’s instructions. The intensities of the bands of SREBP-1c and Lamin A/C in nuclear extracts were quantified with Image analysis software (ImageQuant TL, GE Healthcare). The intensities of nuclear SREBP-1c were normalized by those of corresponding Lamin A/C intensity.

### *In vitro* cleavage assay

Recombinant HSV-human SREBP-1c-HA and human RHBDL4-V5 proteins were prepared using a wheat germ cell-free protein synthesis system with liposomes (CFS) as described previously (Takeda et al., 2015). Proteins were purified using HA and V5-magnetic beads (MBL) respectively and subjected to an *in vitro* cleavage assay as described previously (Urban and Wolfe, 2005). Briefly, purified proteins were reacted in 50 mM Tris-HCl (pH 7.5), 150 mM NaCl, 0.25% DDM buffer for 2 h at 37°C and analyzed by SDS-PAGE and western blot analysis (Takeda et al., 2015).

### Luciferase assay

HEK293 cells were plated on 24-well plates at 5 × 10^4^ cells/well. The following day, the SREBP-1c, SCAP, and RHBDL4 expression plasmids were mixed with SRE-Luc plasmid, or Gal4-VP16-SREBP-1c, RHBDL4, and other expression plasmids indicated in the Figure legends were mixed with Gal4-RE-Luc plasmid. A control plasmid (pCI) to adjust for the total DNA amount per well, and a SV40-*Renilla* luciferase plasmid (pRL-SV40, Promega) as a reference were used. Plasmid DNA and transfection reagents were cotransfected into cultured cells. The next day, luciferase activity in transfectants was measured with Dual-Glo luciferase assay system (Promega) and a fluorometer (Wallac ARVO SX1420 multi-label counter, PerkinElmer). Firefly luciferase activities were normalized to *Renilla* luciferase activities.

### PLAP assay

For the PLAP assay, the next day after cell seeding, PLAP-SREBP-1c, RHBDL4, SCAP plasmids, and a pCI control plasmid to adjust for the total DNA amount per well, were mixed with a firefly luciferase plasmid as reference and cotransfected into cultured cells. After 2 days in culture, the PLAP activity in the culture medium and luciferase activity in the transfectants were measured with the phospha-light system (Applied Biosystems), Dual-Glo luciferase assay system (Promega), and a fluorometer (Wallac ARVO SX1420 multi-label counter, PerkinElmer Japan). PLAP activities were normalized to luciferase activity.

### Gene expression analysis by qRT-PCR

Total RNA was extracted using TRIzol® reagent (Invitrogen), and purified by Purelink Micro-to-Midi total RNA purification system (Invitrogen) according to the manufacturer’s instructions. cDNA was generated using a SuperScript™ III first-strand synthesis system (Invitrogen). Quantitative real-time polymerase chain reaction (qRT-PCR) was performed by using an ABI Prism® 7500 (Applied Biosystems) or Thermal Cycle Dice Real Time System (TP850, Takara Bio) and TB Green Premix Ex Taq II (Takara Bio). Gene expression levels were measured using TaqMan® gene expression assay kit (Applied Biosystems) and the specific primers and TaqMan probes are shown in Table S2. 18s rRNA was measured using TaqMan® ribosomal RNA control reagents (Applied Biosystems) as a control.

### RNA-sequencing analysis

Total RNA was extracted using Sepasol reagent (Nacalai tesque) from the livers of WT and RHBDL4 KO mice (n = 4, each group). RNA-sequencing libraries were prepared with a NEBNext rRNA Depletion Kit and an ENBNext Ultra Directional RNA Library Prep Kit (New England Biolabs). Then, 2 × 36 base paired-end sequencing was performed with a NextSeq500 sequencer (Illumina) by Tsukuba i-Laboratory LLP (Tsukuba, Japan). FASTQ files were analyzed using CLC Genomics Workbench (Version 7.5.1; Qiagen). Sequences were mapped to the mouse genome (mm10) and quantified for annotated genes. Differential expression was analyzed using the Reactome Knowledgebase (https://reactome.org).

### Immunoprecipitation assay

For immunoprecipitation, HEK293A cells were cotransfected with HSV-SREBP-1c, RHBDL4-V5, SCAP-HA, and Myc-Insig1 as described in Figure legends. Cells were lysed with buffer A (10 mM HEPES/NaOH, pH 7.9, 10 mM/L KCl, 1.5 mM MgCl_2_, 1 mM EDTA, 1 mM DTT, 0.1% NP-40, and a protease inhibitor cocktail tablet [Roche]). After clarifying lysates by centrifugation at 2,000 rpm for 1 min at 4 °C twice, supernatant fractions were collected. The supernatant fractions, which were supplemented with KCl to a final concentration of 100 mM were subjected directly to immunoprecipitation and incubated for overnight at 4 °C with antibodies and protein G-Sepharose 4 Fast Flow (GE Healthcare) or HA magnetic beads (Pierce). After repeated washing steps, bound proteins were eluted with SDS sample buffer or HA peptide (MBL) and analyzed by SDS-PAGE and western blot analysis.

### Microscopy

HEK293A cells grown on coverslips were cotransfected with YFP-RHBDL4, mCherry-SREBP-1c, DsRed-ER, YFP-SREBP-1c-CFP, mCherry-SCAP, mCherry-RHBDL4, and mCherry-RHBDL4 (S144A) as described in Figure legends. The cells were fixed with 4 % paraformaldehyde for 15 min and nuclei were visualized by staining with DAPI. Confocal microscopy was performed on a TCS SP8 laser-scanning system (Leica).

### Adenoviral overexpression of RHBDL4

For preparation of recombinant adenovirus, Myc-tagged mouse RHBDL4 and GFP were subcloned into pENTR4 vectors (Invitrogen). Recombinant adenovirus plasmids were homologously recombined with pAd/CMV/V5-DEST vectors (Invitrogen), as per manufacturer’s protocols. Recombinant adenoviruses were produced in HEK293A cells (Invitrogen) and purified by CsCl gradient centrifugation, as described previously (Nakagawa et al., 2006). For adenovirus infection, mice were *i*.*v*. injected with adenovirus at a total dose of 5 × 10^10^ OPU/mouse; total infection amounts of adenovirus were adjusted using GFP adenovirus (Nakagawa et al., 2006). Six days after infection, fasting and refeeding experiments were performed as described previously (Ide et al., 2004).

### Histological analysis and metabolic measurements

Mouse livers were fixed, embedded in paraffin, sectioned, and stained with hematoxylin and eosin (H&E). For Oil red O (ORO) staining, liver tissues were harvested in cold PBS, fixed overnight at 4 °C in 4% paraformaldehyde in PBS, cryoprotected in 30% sucrose in PBS, embedded in optimal cutting temperature compound (Sakura Finetek Inc.), and frozen. Frozen tissues were cut into 5-μm thick cryosections and stained with Oil red O (Sigma-Aldrich). Plasma and liver parameters were measured as described previously (Nakagawa et al., 2006). Fatty acid composition by gas chromatography was measured as described previously (Sekiya et al., 2003).

### Structural modeling of the RHBDL4–SREBP-1c complex structure

The RHBDL4–SREBP-1c complex structure was constructed using the Molecular Operating Environment (MOE) program (2013.08 Chemical Computing Group Inc.: 1010 Sherbrooke St. West, Suite #910, Montreal, QC, Canada, H3A 2R7, 2018). Template structures for homology modeling were taken from the crystal structure of GlpG (PDB ID: 2NRF) (Wu et al., 2006) for RHBDL4 (GenBank accession number: NP_001161080.1) and the structure of the channel-forming transmembrane domain of Virus protein from HIV-1 (PDB ID: 1PJE) (Park et al., 2003) for SREBP-1c (GenBank accession number: NP_001308025.1). Sequence alignment was performed manually using Align utility. The modeled complex structure of the RHBDL4– SREBP-1c was protonated using Protonate 3D module of MOE utilizing the AMBER10:EHT force-field with solvation energy accounted for via the Born model. After the structure was subjected to energy minimization by keeping heavy atoms constrained, the resultant complex structure was fully optimized. The structures were visualized by using PyMOL (The PyMOL Molecular Graphics System, Version 2.0 (Schrodinger, LLC)).

## QUANTIFICATION AND STATISTICAL ANALYSIS

### Statistical analysis

Statistical analysis was performed with Statistical Analysis System Software (SAS Institute, Cary, NC) or GraphPad Prism 8. Comparison between the results of the two groups was performed with the Student’s t-test or nonparametric Wilcoxon matched-pair analyses. Comparisons among the results of multiple groups were made with Dunnett’s multiple comparison tests, the nonparametric Steel test or ANOVA followed by Bonferroni test. Values were considered significant when the p < 0.05. The results were given as the mean ± SEM.

## Supplemental Information titles and legends

**Figure S1.**
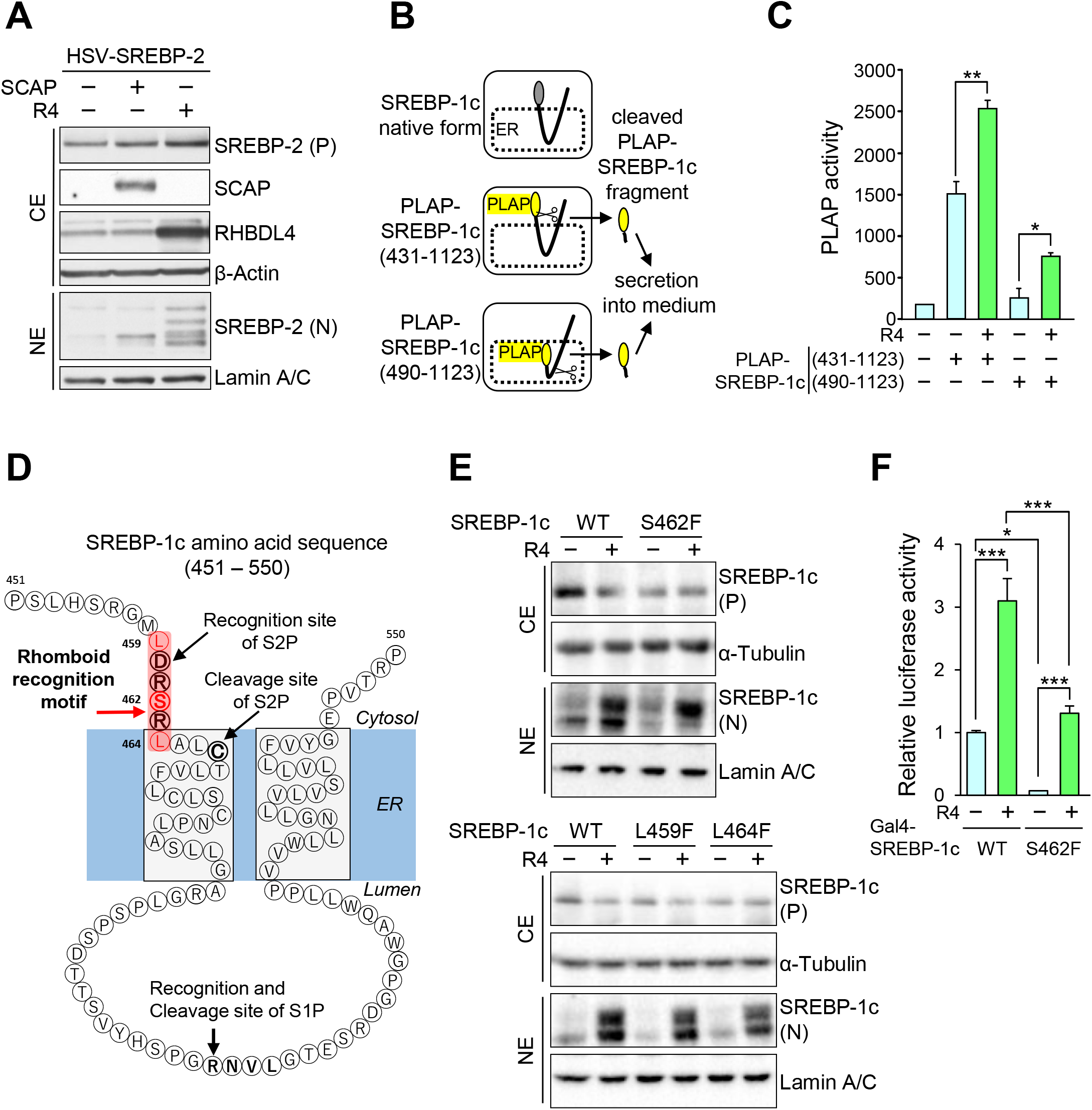
The effect of RHBDL4 on SREBP-2 cleavage and mapping of cleavage region of SREBP-1c, Related to Figure 1. (A) Indicated proteins were expressed in HEK293 cells. After 24 hr, cytosolic extracts (CE) and nuclear extracts (NE) were immunoblotted with indicated antibodies. HSV-SREBPs were immunoblotted with HSV antibodies. P, precursor; N, nuclear. (B and C) Schematic overview of placental alkaline phosphatase (PLAP) assay (B). (C) PLAP-SREBP-1c (431-1123) and PLAP-SREBP-1c (490-1123) were prepared by fusing the catalytic domain of PLAP with the indicated amino acid residues of SREBP-1c. Indicated proteins were expressed in HEK293 cells. After 24 hr, PLAP activities in the medium were measured. Quantification was performed on six samples. (D) Diagram of SREBP-1c (451-550). Transmembrane domains are embedded in gray boxes. Numbers indicate the number of amino acid residues. S1P and S2P recognition and cleavage sites are indicated in bold. Rhomboid recognition motif and mutated amino acids are indicated in pink boxes and red, respectively. (E) Immunoblot analysis of SREBP-1c in CE and NE of HEK293A cells expressing RHBDL4 and either SREBP-1c WT or SREBP-1c harboring the indicated mutants. (F) HEK293A cells were transfected with Gal4-RE-Luc, pRL-SV40, and the indicated expression plasmids. After 24 hr, cell extracts were examined using luciferase assays. Quantification was performed on six samples. Data are represented as means ± SEM. * P < 0.05, ** P < 0.01, *** P < 0.001.

**Figure S2.**
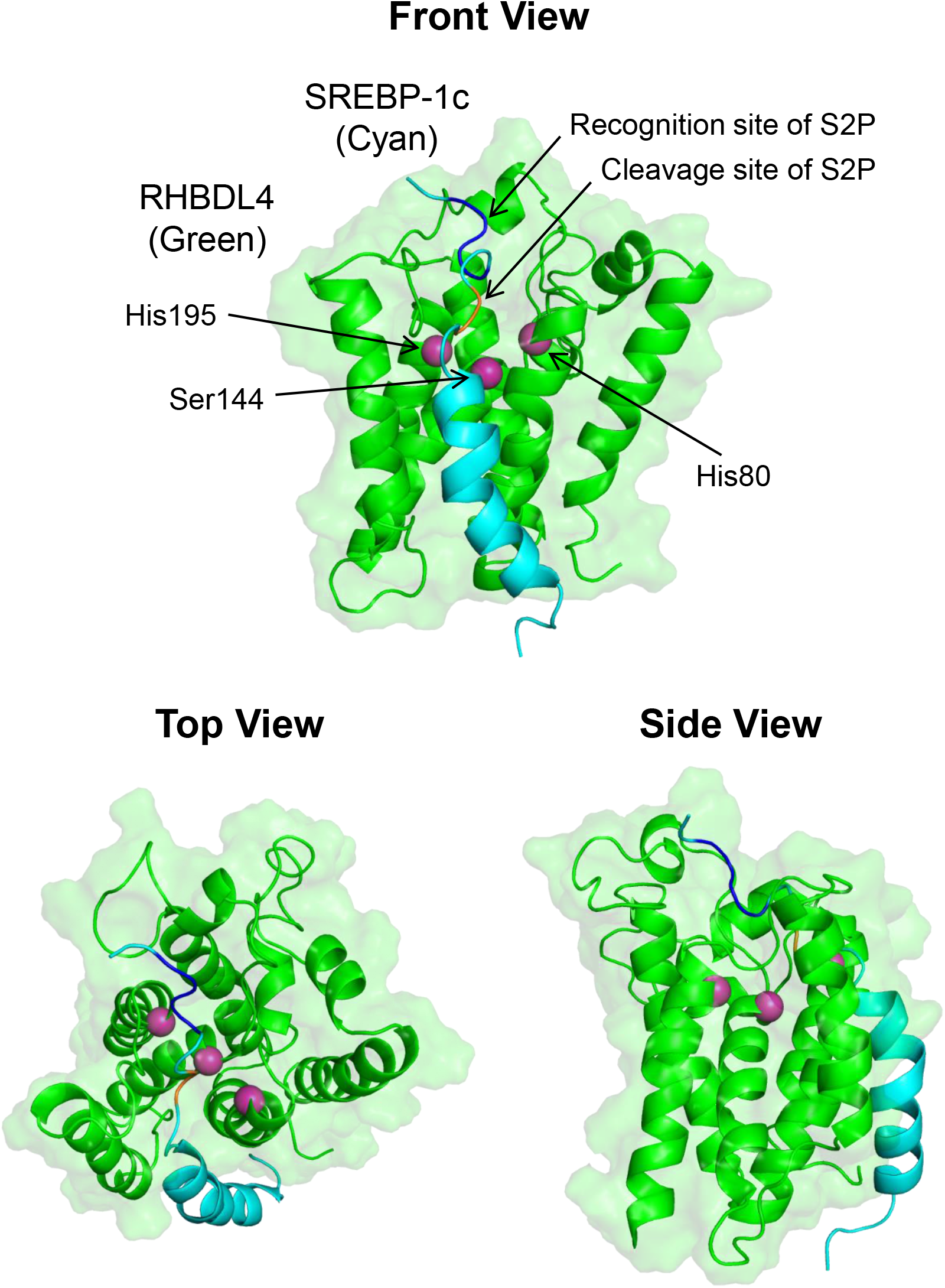
Structure modeling of the RHBDL4–SREBP-1c complex, Related to Figure 1. The RHBDL4 protease core is formed by six helical transmembrane domains (green) and contains a catalytic dyad of serine and histidine. The transmembrane domain of SREBP-1c as the substrate (cyan) interacts with the RHBDL4 active site harboring the catalytic dyad. The S2P cleavage site of SREBP-1c in sterol-regulated classical pathway is located near the catalytic dyad of RHBDL4.

**Figure S3.**
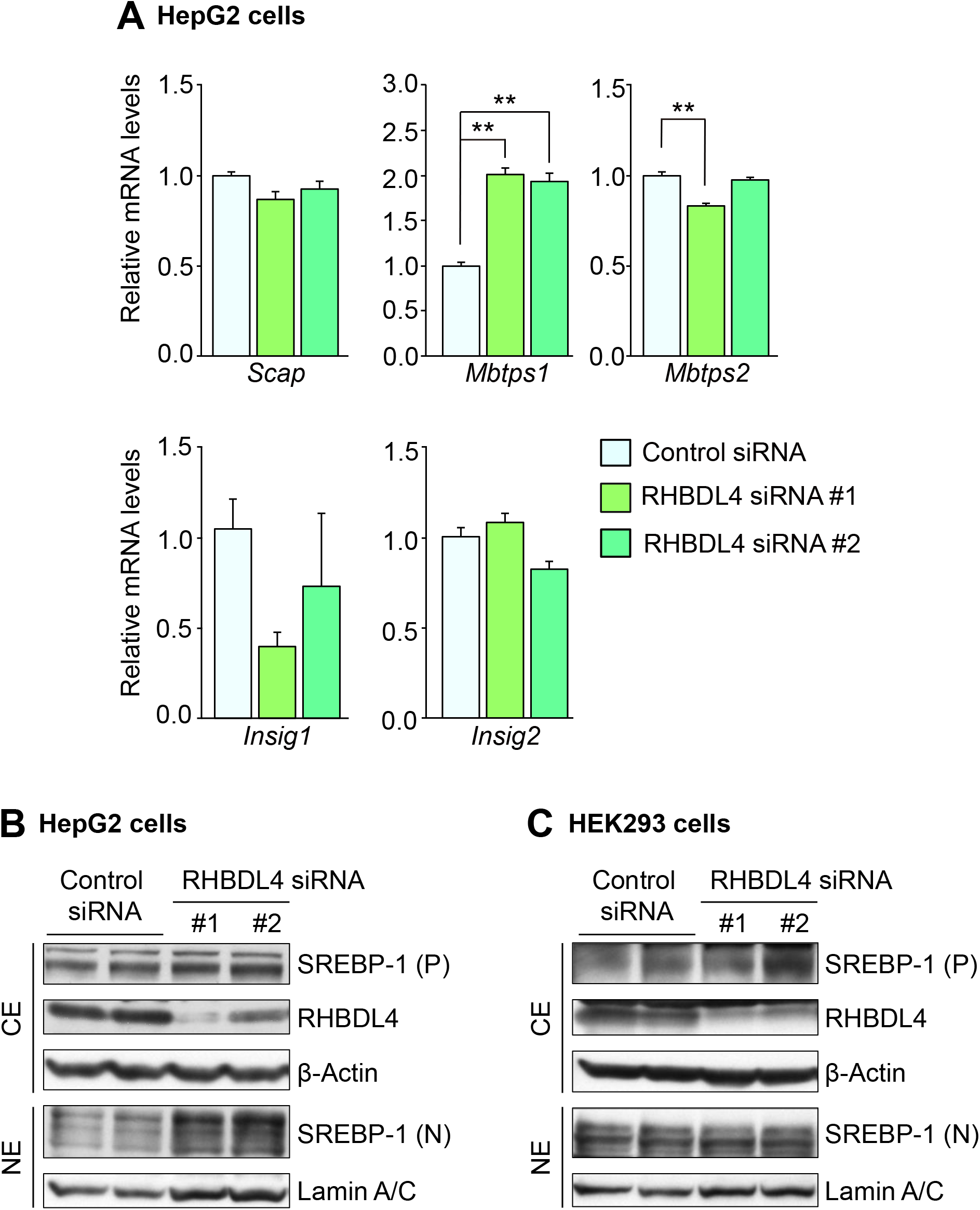
SREBP-1 related gene expression analysis and effect of 25-hydroxycholesterol treatment in siRNA knockdown of RHBDL4 cells, Related to Figure 3. (A) qRT-PCR analysis of SREBP related genes. HepG2 cells were transfected with control siRNA or two independent RHBDL4 siRNAs (#1 and #2). Twenty-four hr after transfection, cells were incubated with 10% DLS containing 3 μM 25-hydroxycholesterol for 24 hr. qRT-PCR analysis of SREBP-1 related genes was performed. Data represented as means ± SEM. Quantification was performed on six samples. ** P < 0.01. (B and C) Immunoblot analysis of the cleavage of SREBP-1 by RHBDL4. HepG2 cells (B) and HEK293 cells (C) were transfected with control siRNA or two independent RHBDL4 siRNAs (#1 and #2). Twenty-four hr after transfection, cells were incubated with 10% DLS for 24 hr. Immunoblot analysis of indicated proteins in cytosolic extracts (CE) and nuclear extracts (NE) was performed. P, precursor; N, nuclear.

**Figure S4.**
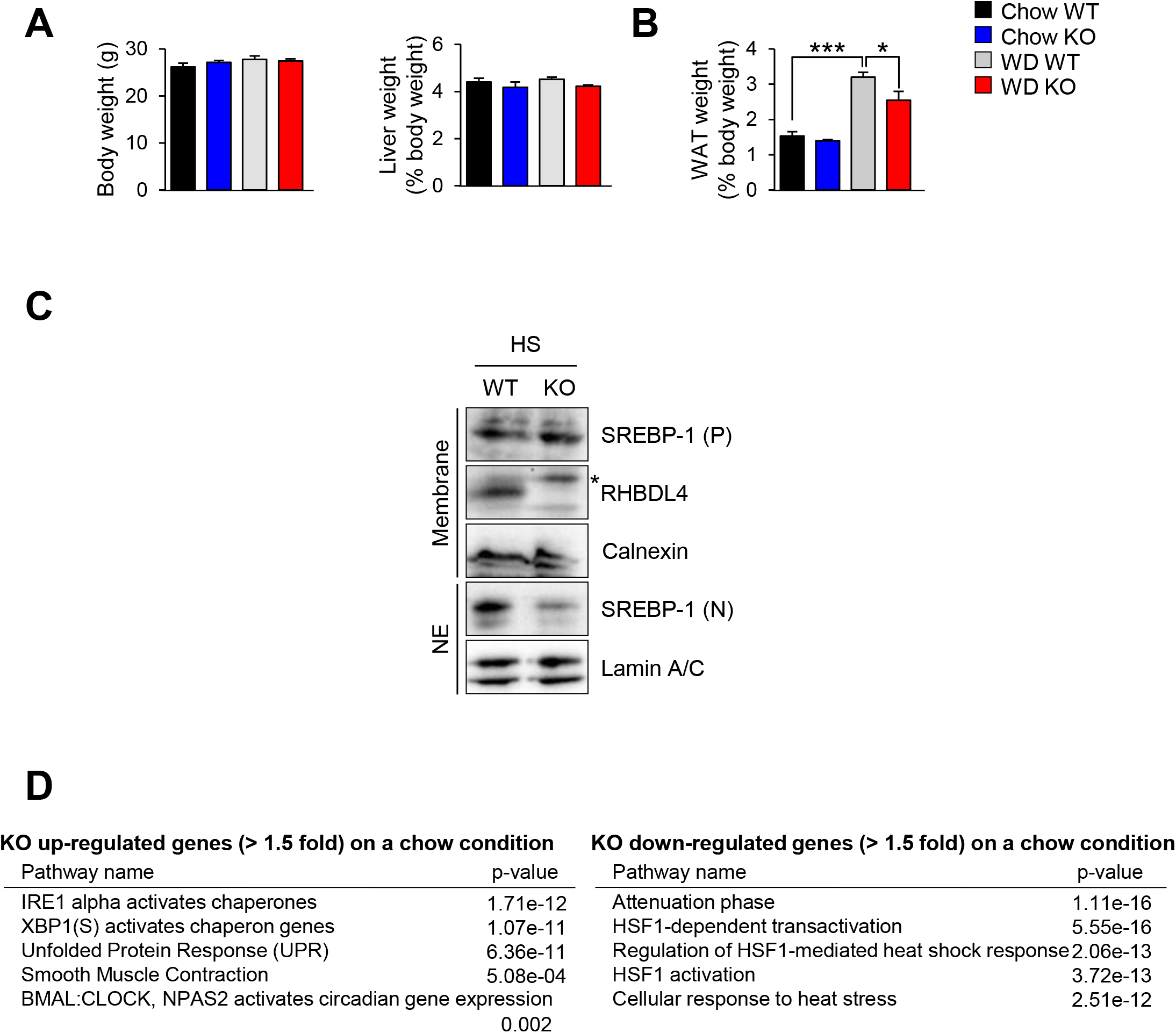
Characteristics of RHBDL4 knockout mice, Related to Figures 4 and 5. Eight-week-old male wild type (WT) and RHBDL4 knockout (KO) mice were fed with normal chow or a Western diet (WD) for 14 days (A, B and D), or a high-sucrose diet (HS) for 7 days (C). (A and B) Body weight and liver weight (A), white adipose tissue (WAT) weight (B) of mice. Data are represented as means ± SEM. Quantification was performed on eight samples (A) or four samples (B). * P < 0.05, *** P < 0.001. (C) Representative immunoblot analysis of the indicated proteins in the membrane fraction and nuclear extracts (NE) of mouse livers. P, precursor; N, nuclear. (D) Functional annotations associated with KO upregulated (left) and downregulated (right) genes compared to WT mice fed a normal chow diet.

**Figure S5.**
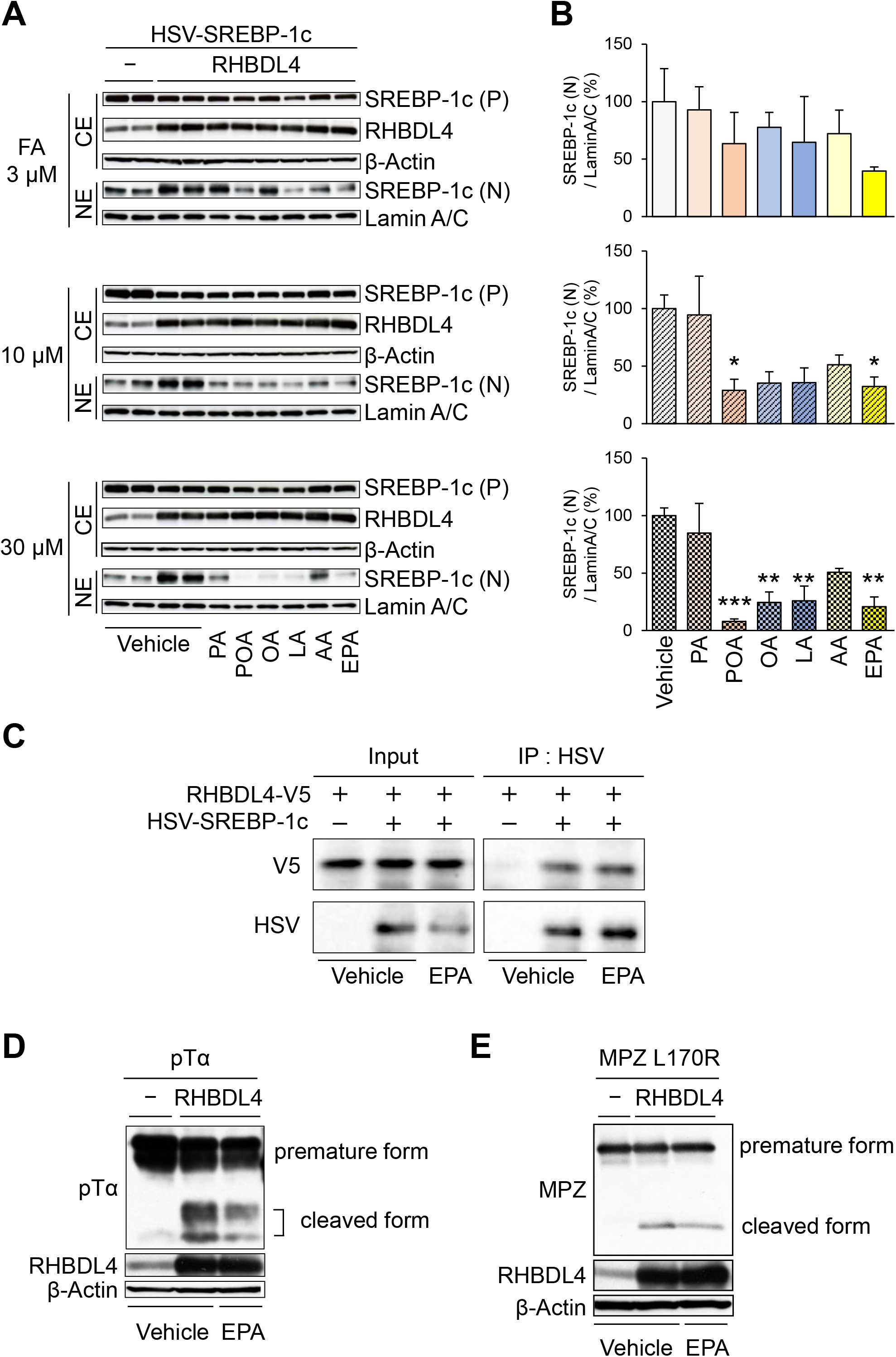
PUFA inhibits and SFA activates RHBDL4-dependent SREBP-1c cleavage, Related to Figure 6. (A) HEK293 cells were transfected with HSV-SREBP-1c and RHBDL4 expression plasmids. After 4 hr, cells were incubated with 10% DLS containing the indicated concentration of each fatty acid for 20 hr. Immunoblot analysis of the indicated proteins in cytosolic extracts (CE) and nuclear extracts (NE) was performed. SREBP-1c cleavage by various fatty acids were shown. P, precursor; N, nuclear. (B) The quantification of the relative levels of SREBP-1c nuclear form to the vehicle treatment of panel (A). Data are represented as means ± SEM from three independent experiments. * P < 0.05, ** P < 0.01, *** P < 0.001 compared with vehicle group. (C) The effect of EPA on the interaction of RHBDL4 and SREBP-1c. HEK293A cells were transfected with HSV-SREBP-1c and RHBDL4-V5 expression plasmids. After 4 hr, cells were incubated with 10% DLS containing with 3 μM EPA for 20 hr. Cell extracts were immunoprecipitated (IP) with anti-HSV. Bound proteins were immunoblotted with anti-HSV and anti-V5 antibodies. (D and E) The suppressive effect of EPA on the cleavage of pTα and MPZ L170R induced by RHBDL4. HEK293 cells were transfected with RHBDL4 and HA-tagged pTα (D) or MPZ L170R (E) expression plasmids. After 4 hr, cells were incubated with 10% DLS containing with 50 μM EPA for 20 hr. Immunoblot analysis of the indicated proteins in cell extracts was performed.

**Figure S6.**
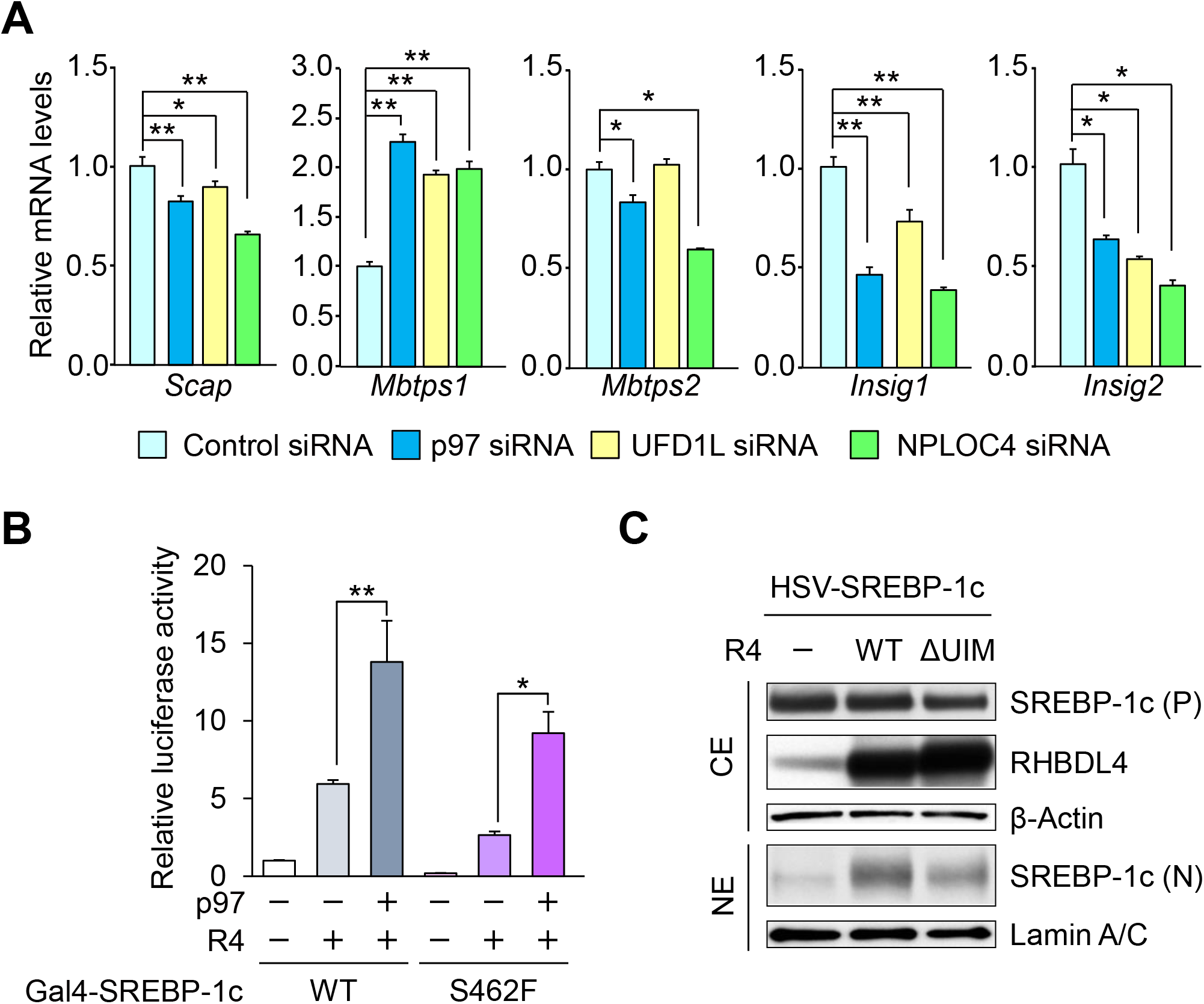
The analysis of gene expression in p97/VCP and related factors through siRNA knockdown and the effect of ubiquitin interaction motif in RHBDL4 on SREBP-1c cleavage, Related to Figure 7. (A) qRT-PCR analysis of SREBP-1 related genes. HepG2 cells were transfected with the indicated siRNA. 24 hr after transfection, cells were incubated with 10% DLS containing 3 μM 25-hydroxycholesterol for 24 hr. Data are represented as means ± SEM. Quantification was performed on six samples. *P < 0.05, ** P < 0.01. (B) Luciferase assay of SREBP-1c regulated by p97. HEK293A cells were transfected with Gal4-RE-Luc, pRL-SV40, and the indicated expression plasmids. After 24 hr, cell extracts were examined using luciferase assays. Data are represented as means ± SEM. Quantification was performed on six samples. *P < 0.05, ** P < 0.01. (C) The RHBDL4 with mutation of ubiquitin interaction motif (UIM) show weaker cleavage activity for SREBP-1c. HEK293 cells were transfected with HSV-SREBP-1c and either RHBDL4 WT or mutant RHBDL4 (ΔUIM) plasmid. The indicated proteins in cytosolic extracts (CE) and nuclear extracts (NE) were immunoblotted. HSV-SREBPs were immunoblotted with HSV antibodies. P, precursor; N, nuclear.

**Table S1.** Fatty acid profile in the liver of WT and RHBDL4 KO mice, Related to Figure 5.

|  | Fatty acid contents (mg/g) |  | Fatty acid composition (molar%) |  |
| --- | --- | --- | --- | --- |
|  | WT | KO | WT | KO |
| C12:0 | 0.006 ± 0.003 | 0.005 ± 0.001 | 0.071 ± 0.030 | 0.073 ± 0.013 |
| C14:0 | 0.055 ± 0.004 | 0.053 ± 0.009 | 0.545 ± 0.026 | 0.703 ± 0.089 |
| C14:1ω5 | 0.006 ± 0.001 | 0.005 ± 0.001 | 0.062 ± 0.004 | 0.080 ± 0.011 |
| C16:0 | 2.976 ± 0.145 | 2.176 ± 0.156** | 26.404 ± 0.185 | 26.344 ± 0.234 |
| C16:1ω7 | 0.388 ± 0.017 | 0.219 ± 0.032** | 3.475 ± 0.064 | 2.637 ± 0.219* |
| C18:0 | 1.560 ± 0.058 | 1.110 ± 0.078** | 12.497 ± 0.226 | 12.130 ± 0.432 |
| C18:1ω9 | 2.113 ± 0.125 | 1.753 ± 0.250 | 17.011 ± 0.561 | 19.120 ± 1.995 |
| C18:2ω6 | 1.372 ± 0.067 | 1.034 ± 0.101* | 11.133 ± 0.232 | 11.506 ± 0.975 |
| C18:3ω6 | 0.026 ± 0.001 | 0.024 ± 0.002 | 0.214 ± 0.007 | 0.263 ± 0.010** |
| C18:3ω3 | 0.010 ± 0.001 | 0.006 ± 0.000* | 0.077 ± 0.007 | 0.068 ± 0.004 |
| C20:0 | 0.027 ± 0.001 | 0.026 ± 0.003 | 0.201 ± 0.010 | 0.262 ± 0.028 |
| C20:1ω9 | 0.039 ± 0.003 | 0.042 ± 0.011 | 0.289 ± 0.015 | 0.412 ± 0.096 |
| C20:2ω6 | 0.018 ± 0.001 | 0.014 ± 0.002 | 0.132 ± 0.004 | 0.144 ± 0.013 |
| C20:3ω9 | 0.107 ± 0.010 | 0.071 ± 0.016 | 0.800 ± 0.075 | 0.708 ± 0.133 |
| C20:3ω6 | 0.426 ± 0.016 | 0.275 ± 0.020** | 3.169 ± 0.045 | 2.786 ± 0.089** |
| C20:4ω6 | 1.730 ± 0.104 | 1.254 ± 0.082* | 12.910 ± 0.166 | 12.823 ± 0.396 |
| C20:5ω3 | 0.079 ± 0.005 | 0.042 ± 0.008** | 0.591 ± 0.026 | 0.431 ± 0.077 |
| C22:0 | 0.080 ± 0.003 | 0.046 ± 0.007** | 0.537 ± 0.032 | 0.422 ± 0.067 |
| C22:1ω9 | 0.003 ± 0.000 | 0.004 ± 0.001 | 0.022 ± 0.001 | 0.033 ± 0.005 |
| C22:4ω6 | 0.034 ± 0.002 | 0.035 ± 0.002 | 0.230 ± 0.002 | 0.328 ± 0.046 |
| C22:5ω3 | 0.043 ± 0.004 | 0.027 ± 0.003* | 0.298 ± 0.013 | 0.256 ± 0.019 |
| C24:0 | 0.045 ± 0.001 | 0.033 ± 0.003** | 0.279 ± 0.016 | 0.275 ± 0.018 |
| C22:6ω3 | 1.260 ± 0.101 | 0.822 ± 0.048** | 8.689 ± 0.288 | 7.816 ± 0.325 |
| C24:1ω9 | 0.058 ± 0.002 | 0.045 ± 0.008 | 0.363 ± 0.017 | 0.382 ± 0.055 |
| Total SFA | 4.750 ± 0.207 | 3.448 ± 0.244** | 40.535 ± 0.382 | 40.209 ± 0.592 |
| Total MUFA | 2.608 ± 0.146 | 2.068 ± 0.285 | 21.223 ± 0.636 | 22.664 ± 2.155 |
| Total PUFA | 5.104 ± 0.297 | 3.605 ± 0.248** | 38.242 ± 0.478 | 37.127 ± 1.570 |
| (DHA+EPA)/AA | 0.771 ± 0.015 | 0.691 ± 0.020* | 0.718 ± 0.014 | 0.643 ± 0.019* |
| Total FFAs | 12.462 ± 0.621 | 9.121 ± 0.659* | 100 | 100 |

Data are represented as means ± SEM.
Quantification was performed on four samples. \* P < 0.05, \*\* P < 0.01.

**Table S2.**
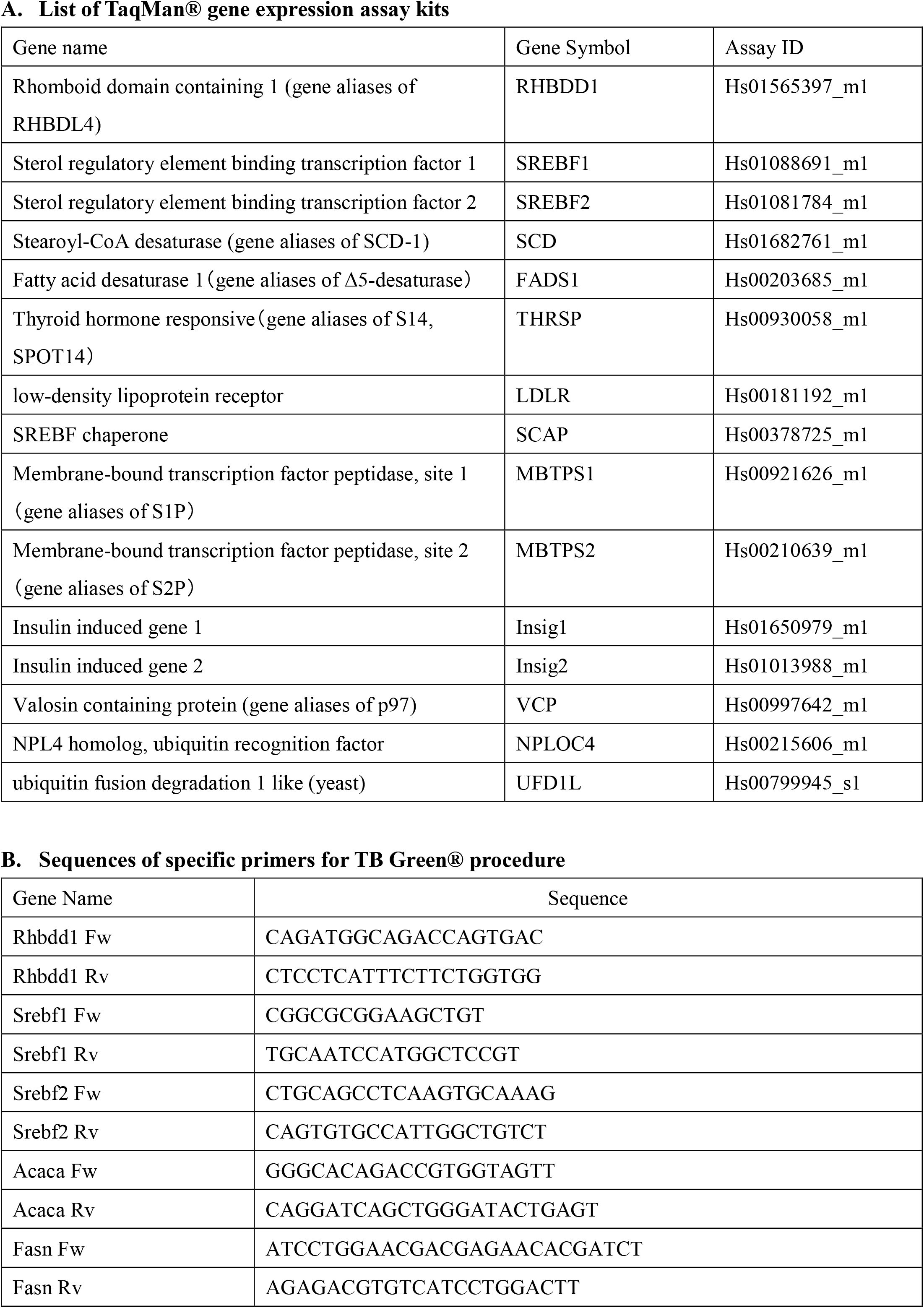
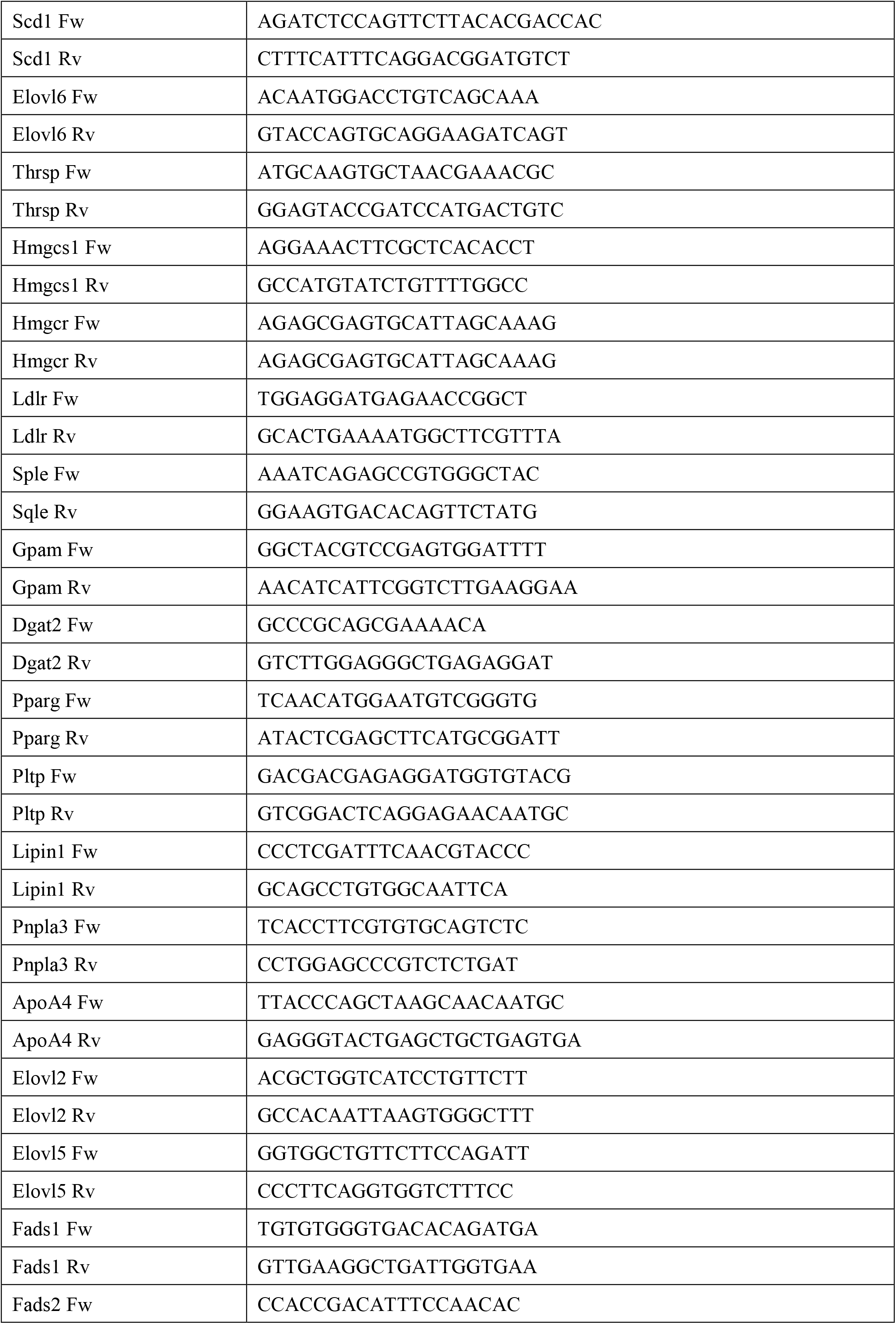
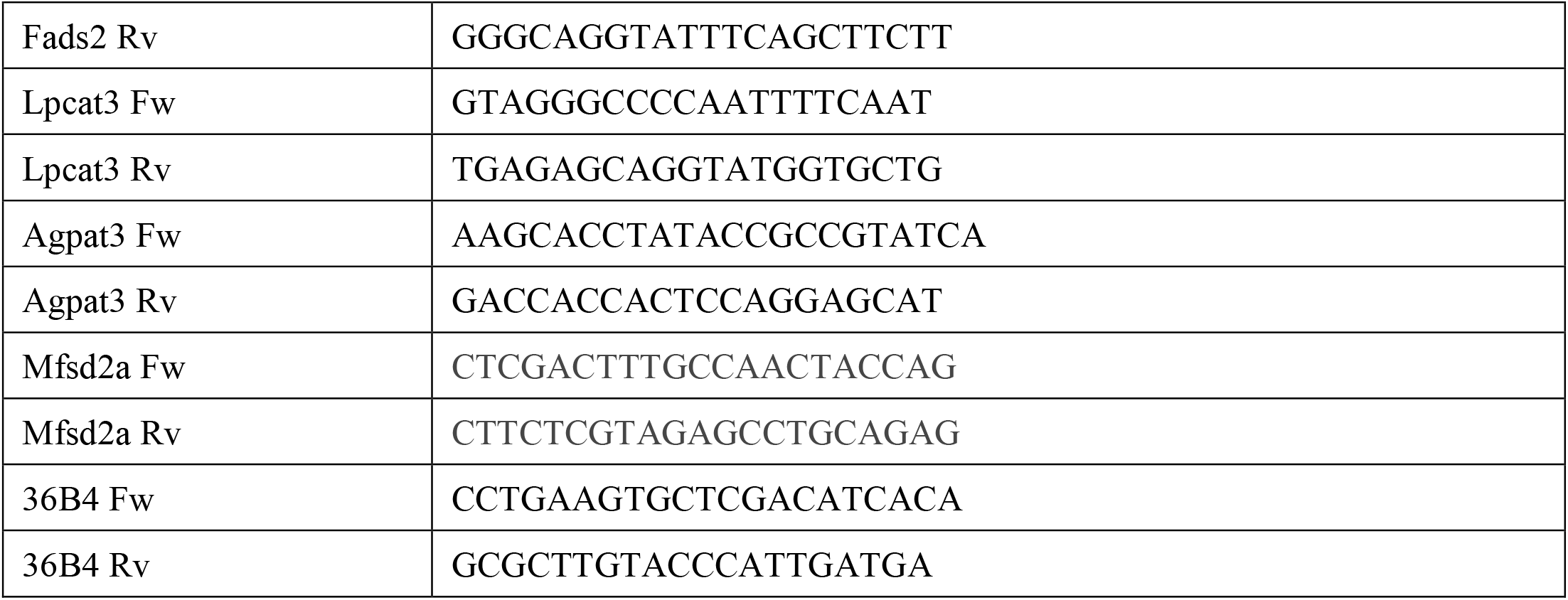
Primers list used in pRT-PCR, Related to Figures 3, 4, 6, 7, S3 and S6.

## REFERENCES

Ben-Zvi, A., Lacoste, B., Kur, E., Andreone, B.J., Mayshar, Y., Yan, H., and Gu, C. (2014). Mfsd2a is critical for the formation and function of the blood-brain barrier. Nature 509, 507–511.

Bergbold, N., and Lemberg, M.K. (2013). Emerging role of rhomboid family proteins in mammalian biology and disease. Biochim Biophys Acta 1828, 2840–2848.

Blythe, E.E., Olson, K.C., Chau, V., and Deshaies, R.J. (2017). Ubiquitin-and ATP-dependent unfoldase activity of P97/VCP*NPLOC4*UFD1L is enhanced by a mutation that causes multisystem proteinopathy. Proc Natl Acad Sci U S A 114, E4380–E4388.

Brown, M.S., and Goldstein, J.L. (1999). A proteolytic pathway that controls the cholesterol content of membranes, cells, and blood. Proc Natl Acad Sci U S A 96, 11041–11048.

Chan, J.P., Wong, B.H., Chin, C.F., Galam, D.L.A., Foo, J.C., Wong, L.C., Ghosh, S., Wenk, M.R., Cazenave-Gassiot, A., and Silver, D.L. (2018). The lysolipid transporter Mfsd2a regulates lipogenesis in the developing brain. PLoS Biol 16, e2006443.

Cong, L., Ran, F.A., Cox, D., Lin, S., Barretto, R., Habib, N., Hsu, P.D., Wu, X., Jiang, W., Marraffini, L.A., et al. (2013). Multiplex genome engineering using CRISPR/Cas systems. Science 339, 819–823.

Fleig, L., Bergbold, N., Sahasrabudhe, P., Geiger, B., Kaltak, L., and Lemberg, M.K. (2012). Ubiquitin-dependent intramembrane rhomboid protease promotes ERAD of membrane proteins. Mol Cell 47, 558–569.

Gordon, J.W., and Ruddle, F.H. (1981). Integration and stable germ line transmission of genes injected into mouse pronuclei. Science 214, 1244–1246.

Greenblatt, E.J., Olzmann, J.A., and Kopito, R.R. (2012). Making the cut: intramembrane cleavage by a rhomboid protease promotes ERAD. Nat Struct Mol Biol 19, 979–981.

Hannah, V.C., Ou, J., Luong, A., Goldstein, J.L., and Brown, M.S. (2001). Unsaturated fatty acids down-regulate srebp isoforms 1a and 1c by two mechanisms in HEK-293 cells. J Biol Chem 276, 4365–4372.

Hashidate-Yoshida, T., Harayama, T., Hishikawa, D., Morimoto, R., Hamano, F., Tokuoka, S.M., Eto, M., Tamura-Nakano, M., Yanobu-Takanashi, R., Mukumoto, Y., et al. (2015). Fatty acid remodeling by LPCAT3 enriches arachidonate in phospholipid membranes and regulates triglyceride transport. Elife 4.

Hishikawa, D., Yanagida, K., Nagata, K., Kanatani, A., Iizuka, Y., Hamano, F., Yasuda, M., Okamura, T., Shindou, H., and Shimizu, T. (2020). Hepatic Levels of DHA-Containing Phospholipids Instruct SREBP1-Mediated Synthesis and Systemic Delivery of Polyunsaturated Fatty Acids. iScience 23, 101495.

Horton, J.D., Goldstein, J.L., and Brown, M.S. (2002). SREBPs: activators of the complete program of cholesterol and fatty acid synthesis in the liver. J Clin Invest 109, 1125–1131.

Horton, J.D., Shimomura, I., Brown, M.S., Hammer, R.E., Goldstein, J.L., and Shimano, H. (1998). Activation of cholesterol synthesis in preference to fatty acid synthesis in liver and adipose tissue of transgenic mice overproducing sterol regulatory element-binding protein-2. J Clin Invest 101, 2331–2339.

Hwang, J., Ribbens, D., Raychaudhuri, S., Cairns, L., Gu, H., Frost, A., Urban, S., and Espenshade, P.J. (2016). A Golgi rhomboid protease Rbd2 recruits Cdc48 to cleave yeast SREBP. EMBO J 35, 2332–2349.

Ide, T., Shimano, H., Yahagi, N., Matsuzaka, T., Nakakuki, M., Yamamoto, T., Nakagawa, Y., Takahashi, A., Suzuki, H., Sone, H., et al. (2004). SREBPs suppress IRS-2-mediated insulin signalling in the liver. Nat Cell Biol 6, 351–357.

Jeon, T.I., and Osborne, T.F. (2012). SREBPs: metabolic integrators in physiology and metabolism. Trends Endocrinol Metab 23, 65–72.

Kim, J., Ha, H.J., Kim, S., Choi, A.R., Lee, S.J., Hoe, K.L., and Kim, D.U. (2015). Identification of Rbd2 as a candidate protease for sterol regulatory element binding protein (SREBP) cleavage in fission yeast. Biochem Biophys Res Commun 468, 606–610.

Kreutzberger, A.J.B., Ji, M., Aaron, J., Mihaljevic, L., and Urban, S. (2019). Rhomboid distorts lipids to break the viscosity-imposed speed limit of membrane diffusion. Science 363.

Lee, J.N., Zhang, X., Feramisco, J.D., Gong, Y., and Ye, J. (2008). Unsaturated fatty acids inhibit proteasomal degradation of Insig-1 at a postubiquitination step. J Biol Chem 283, 33772–33783.

Lemberg, M.K. (2011). Intramembrane proteolysis in regulated protein trafficking. Traffic 12, 1109–1118.

Moin, S.M., and Urban, S. (2012). Membrane immersion allows rhomboid proteases to achieve specificity by reading transmembrane segment dynamics. Elife 1, e00173.

Moon, Y.A., Hammer, R.E., and Horton, J.D. (2009). Deletion of ELOVL5 leads to fatty liver through activation of SREBP-1c in mice. J Lipid Res 50, 412–423.

Nakagawa, Y., Shimano, H., Yoshikawa, T., Ide, T., Tamura, M., Furusawa, M., Yamamoto, T., Inoue, N., Matsuzaka, T., Takahashi, A., et al. (2006). TFE3 transcriptionally activates hepatic IRS-2, participates in insulin signaling and ameliorates diabetes. Nat Med 12, 107–113.

Nakakuki, M., Kawano, H., Notsu, T., Imada, K., Mizuguchi, K., and Shimano, H. (2014). A novel processing system of sterol regulatory element-binding protein-1c regulated by polyunsaturated fatty acid. J Biochem 155, 301–313.

Nguyen, L.N., Ma, D., Shui, G., Wong, P., Cazenave-Gassiot, A., Zhang, X., Wenk, M.R., Goh, E.L., and Silver, D.L. (2014). Mfsd2a is a transporter for the essential omega-3 fatty acid docosahexaenoic acid. Nature 509, 503–506.

Olzmann, J.A., Richter, C.M., and Kopito, R.R. (2013). Spatial regulation of UBXD8 and p97/VCP controls ATGL-mediated lipid droplet turnover. Proc Natl Acad Sci U S A 110, 1345–1350.

Park, S.H., Mrse, A.A., Nevzorov, A.A., Mesleh, M.F., Oblatt-Montal, M., Montal, M., and Opella, S.J. (2003). Three-dimensional structure of the channel-forming trans-membrane domain of virus protein “u” (Vpu) from HIV-1. J Mol Biol 333, 409–424.

Pauter, A.M., Olsson, P., Asadi, A., Herslof, B., Csikasz, R.I., Zadravec, D., and Jacobsson, A. (2014). Elovl2 ablation demonstrates that systemic DHA is endogenously produced and is essential for lipid homeostasis in mice. J Lipid Res 55, 718–728.

Radhakrishnan, S.K., den Besten, W., and Deshaies, R.J. (2014). p97-dependent retrotranslocation and proteolytic processing govern formation of active Nrf1 upon proteasome inhibition. Elife 3, e01856.

Rape, M., Hoppe, T., Gorr, I., Kalocay, M., Richly, H., and Jentsch, S. (2001). Mobilization of processed, membrane-tethered SPT23 transcription factor by CDC48(UFD1/NPL4), a ubiquitin-selective chaperone. Cell 107, 667–677.

Rong, X., Wang, B., Palladino, E.N., de Aguiar Vallim, T.Q., Ford, D.A., and Tontonoz, P. (2017). ER phospholipid composition modulates lipogenesis during feeding and in obesity. J Clin Invest 127, 3640–3651.

Sato, A., Kawano, H., Notsu, T., Ohta, M., Nakakuki, M., Mizuguchi, K., Itoh, M., Suganami, T., and Ogawa, Y. (2010). Antiobesity effect of eicosapentaenoic acid in high-fat/high-sucrose diet-induced obesity: importance of hepatic lipogenesis. Diabetes 59, 2495–2504.

Sekiya, M., Yahagi, N., Matsuzaka, T., Najima, Y., Nakakuki, M., Nagai, R., Ishibashi, S., Osuga, J., Yamada, N., and Shimano, H. (2003). Polyunsaturated fatty acids ameliorate hepatic steatosis in obese mice by SREBP-1 suppression. Hepatology 38, 1529–1539.

Sekiya, M., Yahagi, N., Matsuzaka, T., Takeuchi, Y., Nakagawa, Y., Takahashi, H., Okazaki, H., Iizuka, Y., Ohashi, K., Gotoda, T., et al. (2007). SREBP-1-independent regulation of lipogenic gene expression in adipocytes. J Lipid Res 48, 1581–1591.

Sheng, Z., Otani, H., Brown, M.S., and Goldstein, J.L. (1995). Independent regulation of sterol regulatory element-binding proteins 1 and 2 in hamster liver. Proc Natl Acad Sci U S A 92, 935–938.

Shimano, H., Horton, J.D., Shimomura, I., Hammer, R.E., Brown, M.S., and Goldstein, J.L. (1997). Isoform 1c of sterol regulatory element binding protein is less active than isoform 1a in livers of transgenic mice and in cultured cells. J Clin Invest 99, 846–854.

Shimano, H., and Sato, R. (2017). SREBP-regulated lipid metabolism: convergent physiology - divergent pathophysiology. Nat Rev Endocrinol 13, 710–730.

Shimomura, I., Shimano, H., Horton, J.D., Goldstein, J.L., and Brown, M.S. (1997). Differential expression of exons 1a and 1c in mRNAs for sterol regulatory element binding protein-1 in human and mouse organs and cultured cells. J Clin Invest 99, 838–845.

Singh, A.N., Oehler, J., Torrecilla, I., Kilgas, S., Li, S., Vaz, B., Guerillon, C., Fielden, J., Hernandez-Carralero, E., Cabrera, E., et al. (2019). The p97-Ataxin 3 complex regulates homeostasis of the DNA damage response E3 ubiquitin ligase RNF8. EMBO J 38, e102361.

Strisovsky, K., Sharpe, H.J., and Freeman, M. (2009). Sequence-specific intramembrane proteolysis: identification of a recognition motif in rhomboid substrates. Mol Cell 36, 1048–1059.

Takeda, H., Ogasawara, T., Ozawa, T., Muraguchi, A., Jih, P.J., Morishita, R., Uchigashima, M., Watanabe, M., Fujimoto, T., Iwasaki, T., et al. (2015). Production of monoclonal antibodies against GPCR using cell-free synthesized GPCR antigen and biotinylated liposome-based interaction assay. Sci Rep 5, 11333.

Takeuchi, Y., Yahagi, N., Izumida, Y., Nishi, M., Kubota, M., Teraoka, Y., Yamamoto, T., Matsuzaka, T., Nakagawa, Y., Sekiya, M., et al. (2010). Polyunsaturated fatty acids selectively suppress sterol regulatory element-binding protein-1 through proteolytic processing and autoloop regulatory circuit. J Biol Chem 285, 11681–11691.

Torrecilla, I., Oehler, J., and Ramadan, K. (2017). The role of ubiquitin-dependent segregase p97 (VCP or Cdc48) in chromatin dynamics after DNA double strand breaks. Philos Trans R Soc Lond B Biol Sci 372.

Urban, S., and Wolfe, M.S. (2005). Reconstitution of intramembrane proteolysis in vitro reveals that pure rhomboid is sufficient for catalysis and specificity. Proc Natl Acad Sci U S A 102, 1883–1888.

Varin, A., Thomas, C., Ishibashi, M., Menegaut, L., Gautier, T., Trousson, A., Bergas, V., de Barros, J.P., Narce, M., Lobaccaro, J.M., et al. (2015). Liver X receptor activation promotes polyunsaturated fatty acid synthesis in macrophages: relevance in the context of atherosclerosis. Arterioscler Thromb Vasc Biol 35, 1357–1365.

VerHague, M.A., Cheng, D., Weinberg, R.B., and Shelness, G.S. (2013). Apolipoprotein A-IV expression in mouse liver enhances triglyceride secretion and reduces hepatic lipid content by promoting very low density lipoprotein particle expansion. Arterioscler Thromb Vasc Biol 33, 2501–2508.

Wu, Z., Yan, N., Feng, L., Oberstein, A., Yan, H., Baker, R.P., Gu, L., Jeffrey, P.D., Urban, S., and Shi, Y. (2006). Structural analysis of a rhomboid family intramembrane protease reveals a gating mechanism for substrate entry. Nat Struct Mol Biol 13, 1084–1091.

Wunderle, L., Knopf, J.D., Kuhnle, N., Morle, A., Hehn, B., Adrain, C., Strisovsky, K., Freeman, M., and Lemberg, M.K. (2016). Rhomboid intramembrane protease RHBDL4 triggers ER-export and non-canonical secretion of membrane-anchored TGFalpha. Sci Rep 6, 27342.

Xia, D., Tang, W.K., and Ye, Y. (2016). Structure and function of the AAA+ ATPase p97/Cdc48p. Gene 583, 64–77.

Yahagi, N., Shimano, H., Hasty, A.H., Amemiya-Kudo, M., Okazaki, H., Tamura, Y., Iizuka, Y., Shionoiri, F., Ohashi, K., Osuga, J., et al. (1999). A crucial role of sterol regulatory element-binding protein-1 in the regulation of lipogenic gene expression by polyunsaturated fatty acids. J Biol Chem 274, 35840–35844.

